# PhyRepID: a comparative phylogenomics approach for large-scale quantification of protein repeat evolution

**DOI:** 10.1101/2020.02.14.947036

**Authors:** I.A.E.M. van Belzen, E. S. Deutekom, B. Snel

## Abstract

Protein repeats consisting of domains or motifs are involved in key biological processes such as neural development, host-pathogen interactions, and speciation. Expansion and contraction of these repeats can strongly impact protein function as was shown for KNL1 and PRDM9. However, these known cases could only be identified manually and were previously incorrectly reported as conserved in large-scale analyses, because signatures of repeat evolution are difficult to resolve automatically.

We developed PhyRepID to compare protein domain repeat evolution and analysed 4939 groups of orthologous proteins (OGs) from 14 vertebrate species. Our main contributions are 1) detecting a wide scope of repeats consisting of Pfam structural domains and motifs, 2) improving sensitivity and precision of repeat unit detection through optimization for the OGs, 3) using phylogenetic analysis to detect evolution within repeat regions. From these phylogenetic signals, we derived a “protein repeat duplication” (PRD) score that quantifies evolution in repeat regions and thereby enables large-scale comparison of protein families. Zinc finger repeats show remarkably fast evolution, comprising 25 of 100 fastest evolving proteins in our dataset, whilst cooperatively-folding domain repeats like beta-propellers are mostly conserved. Motif repeats have a similar PRD score distribution as domain repeats and also show a large diversity in evolutionary rates. A ranking based on the PRD score reflects previous manual observations of both highly conserved (CDC20) and rapidly evolving repeats (KNL1, PRDM9) and proposes novel candidates (e.g. AHNAK, PRX, SPATA31) showing previously undescribed rapid repeat evolution. PhyRepID is available on https://github.com/ivanbelzen/PhyRepID/.

## Introduction

Protein repeat evolution is a source of variation and a potential mechanism for the evolution of novel adaptive changes (Andrade et al. 2001). Repeat proteins are versatile and involved in many biological processes e.g. cellular structure and transcription. The evolutionary flexibility of repeats contributes to this versatility because gain and loss of existing repeat units is easier than *de novo* formation of energetically stable structures (Andrade et al. 2001). Numerous biologically interesting examples of rapid protein repeat evolution were characterized via manual analyses, e.g. in protein families associated with neural development, host-pathogen interactions (Schüler and Bornberg-Bauer 2016), speciation events (Schwartz et al. 2014; Tromer et al. 2015), human disease (Hanson and Hollingsworth 2016; Yeon et al. 2017) and repression of transposons (Jacobs et al. 2014; Yang et al. 2017). For example, recombination regulator PRDM9 has a rapidly evolving zinc finger array (Schaper et al. 2014; Schüler and Bornberg-Bauer 2016) and has been implicated in (hybrid) sterility and speciation (Schwartz et al. 2014). Kinetochore scaffold protein KNL1/CASC5 has a conserved function in chromosome segregation but shows substantial variation in its binding motif repeat region which is associated with speciation events (Tromer et al. 2015; Roy et al. 2020).

Repeats in proteins consist of similar (or homologous) amino acid subsequences that mostly occur in tandem but also interspersed throughout the protein. The repeating subsequence is referred to as the “repeat unit” and the part of the proteins covered by repeats is the repeat region. About 25% of the human proteome is estimated to contain such repeats (Pellegrini 2015). Repeat units can be either domains or motif sequences (hereafter motifs). Domains have a well-defined structure and function, and consequently are often biophysically constrained in copy number, e.g. if they require cooperative folding (Kajava 2012). Domain databases such as SMART, CDD or Pfam provide profiles for most well-described repeating domains (Punta et al. 2012), e.g. beta-propeller, tetratricopeptide repeat (TPR), leucine-rich repeat (LRR) and ankaryn (Ank). In contrast, motifs lack a defined fold and have generally shorter repeat units than domains. In some resources, short linear motifs are defined as 3-15 amino acids (aa) (Davey et al. 2011), but we detected motifs in a broader range of 10-30 amino acids to enable subsequent alignment. Few motifs have a known function, although some contain one or more kinase target sites or other recognition sites for protein interactions (Davey et al. 2011). Previous work that quantified protein repeat evolution ignored motifs and primarily relied on default detection provided by domain databases for repeat unit identification (Schaper et al. 2014; Persi et al. 2016; Schüler and Bornberg-Bauer 2016).

Protein repeat evolution results in variation in repeat unit copy number, identity and/or ordering between otherwise orthologous proteins. Duplication and loss of repeat units can be caused by replication errors during homologous recombination (Björklund et al. 2006). The resulting repeats within an orthologous group (OG) are challenging to analyse with conventional bioinformatic methods such as multiple sequence alignment, especially since they show varying sequence similarity and divergence from the ancestral repeat region (Andrade et al. 2001). Additionally, it has proven to be a challenge to capture these evolutionary dynamics in large-scale analyses and well-documented rapidly evolving cases are often missed (Schaper et al. 2014; Persi et al. 2016; Schüler and Bornberg-Bauer 2016). Consequently, the fast evolution of protein repeats has been concluded to rarely occur compared to the readily observable rapid expansion and contraction of DNA repeats (Guy-Franck Richard 2000; Schaper et al. 2014; Schüler and Bornberg-Bauer 2016)

We propose a novel approach to compare repeat region evolution between protein families in a quantitative way. We developed the PhyRepID pipeline that uses Phylogenomics to perform Repeat IDentification. First, repeats of domains and motifs are detected using Hidden Markov Models (HMMs) that are optimized for each protein family to attain high sensitivity and precision. Using the homology between the repeat units, phylogenetic trees of the repeats are constructed. Second, evolutionary events, i.e. duplication and loss, are inferred by comparing repeat trees to gene trees using tree reconciliation. The repeat duplications are mapped to the gene tree to indicate where/when the evolutionary events have taken place. Third, this annotated gene tree is reconciled with the species tree to enable comparison of repeat region evolution between OGs. Hereto, the protein repeat duplication (PRD) score is calculated using the number of post-ancestral duplications and the number of proteins in the OG. We show that this PRD score can be used as a relative measure of repeat evolution which describes a gradient of repeat evolutionary dynamics. We find protein domains with structural biophysical constraints, e.g. beta propellers, LRR, TPR and Ank are more often conserved as expected while we find an overrepresentation of C2H2 zinc fingers OGs amongst the fastest 100 OGs (9.2x enrichment). The scoring identified known rapidly evolving protein repeat families (e.g. PRDM9, KNL1) amongst the fastest in our dataset, as well as provide novel candidates for which rapid repeat evolution has not been described before.

## New Approaches

The PhyRepID pipeline (https://github.com/ivanbelzen/PhyRepID/) quantifies repeat evolution using comparative (phylo)genomics of orthologous groups (OGs). The pipeline consists of three components: 1) detection of a broad spectrum of protein repeats consisting of either structural domains or motif sequences, 2) improving the sensitivity and precision of detection by making OG-specific repeat unit HMMs, 3) inferring evolutionary events in the repeat region through phylogenetic comparison of repeat trees to gene trees. Finally, to quantify repeat evolution and compare protein families, a PRD score is derived from the post-ancestral duplications and the number of proteins in the OG (Supplementary Table 1).

### Identifying proteins with repeats consisting of domains or motifs

Prior to repeat detection, protein sequences and orthology were acquired from ENSEMBL (Zerbino et al. 2018) from 14 representative vertebrate species. OGs were filtered on containing at least one human protein-coding gene. We identified 2263 OGs with domain repeats of three or more repeat units in a human protein (Figure 1). HMMs from the Pfam-A database were used for initial domain detection (Finn et al. 2016). Characterization on the level of homologous models (Pfam clans) shows domains known to form tandem repeats comprise the majority (Björklund et al. 2006). The top five Pfam clans in our dataset are beta-propeller (5.7%), C2HC zinc finger (4.6%), TPR (3.7%), LRR (3.0%) and Ank (2.5%) (Table 1). Since not all known repeats are captured by Pfam (e.g. repeat region with binding sites in KNL1), we used MEME for *de novo* detection of motif sequences of 10-30 amino acids (Bailey and Elkan 1994). Motif repeat regions were detected in additional 2676 OGs without domain repeats, after excluding sequence matching known Pfam domains (Figure 1c). To ensure unbiased downstream analyses of the repeat types, the domain and motif datasets were combined to a total of 4939 OGs of which 54% motifs and 46% Pfam domains.

**Figure 1:**
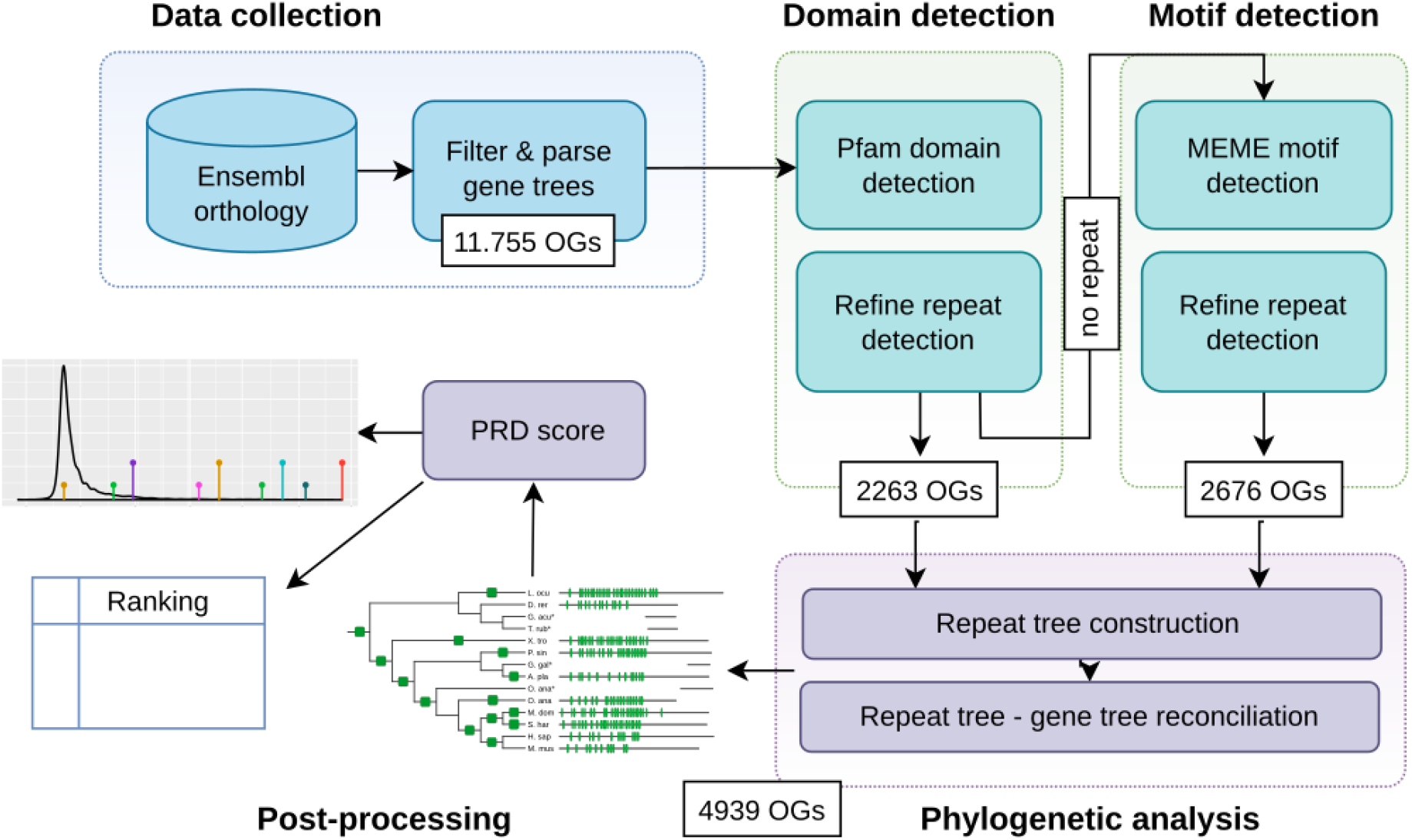
Overview of the PhyRepID pipeline. **Data collection:** Human protein-coding genes were acquired from ENSEMBL, as well as their orthologs in 13 other vertebrate species forming 11.755 orthologous groups (OGs). **Domain detection:** HMMs from Pfam were used to search domains. OG-specific HMMs were made from the best-matching domains to optimize detection of repeat units. Filtering for >=3 repeat units in a human protein resulted in 2363 OGs with domain repeats. **Motif detection:** For OGs where no domain repeat was detected, MEME was used to search for de novo motifs after masking sequence matching to Pfam domains. Following the same procedure as for domain repeats: OG-specific HMMs were made and after filtering this resulted in 2676 OGs with motif repeats. **Phylogenetic analysis:** Phylogenetic trees were made using IQ-TREE for all 4939 OGs from their repeat units and these repeat trees were reconciled with gene trees using TreeFix to infer duplications and losses. **Post-processing:** Downstream analysis was done using various R and python scripts. For each OG a PRD score was calculated with which a ranking was made to compare OGs to each other and find OGs with rapidly-evolving repeats.

**Table 1:** Characteristics of the most prevalent repeat unit classes. #OGs: number of orthologous groups in the dataset, fraction: #OGs/total dataset size. PRD score: the mean PRD scores, all distributions are significantly different from the other groups (less/greater with Wilcoxon test p<0.01. Note: full dataset mean PRD score: 0.01). Post-MRCA duplications: mean repeat duplication count inferred after after the gene tree root. Gene duplications: mean duplication count as annotated on the ENSEMBL Compara gene trees.

| Clan | #OGs | Fraction | PRD score | Post-MRCA duplications | Gene duplications |
| --- | --- | --- | --- | --- | --- |
| motif | 2676 | 0.54 | 0.03 | 2.23 | 1.65 |
| beta propeller | 280 | 0.06 | -0.09 | 0.99 | 1.45 |
| C2H2-zf | 225 | 0.05 | 0.38 | 10.11 | 6.96 |
| TPR | 181 | 0.04 | -0.11 | 0.77 | 1.35 |
| LRR | 148 | 0.03 | -0.12 | 0.70 | 1.11 |
| Ank | 125 | 0.03 | -0.10 | 0.83 | 2.43 |

### Sensitive repeat identification with OG-specific models

In order to prevent spurious repeat dynamics, we aimed to identify repeats with high sensitivity and precision. Repeat unit detection was refined by optimizing a profile HMM specific for the OG through an iterative process (Figure 1). This circumvents detection issues (i.e. missing or partial units) due to sequence divergence between protein families (Stirnimann et al. 2010; Punta et al. 2012). Prior to optimization, Pfam domain models are grouped by homology according to their Pfam clans, which resolves issues of within-protein overlap and improves interpretation of biologically relevant differences within and between species. This refinement process improved the sensitivity and precision of domain detection, reducing the differences detected between orthologs compared to the default annotation for 51% (1152) of the Pfam OGs as shown by a decrease in the coefficient of variation in the repeat unit count (Supplementary Table 2).

Notably: variation in repeat unit count disappeared in 199 Pfam OGs (9%), highlighting the importance of sensitive and precise detection for preventing inference of false positive evolutionary events. After the refinement process, Pfam domains maintain their characteristic length and number of units (Björklund et al. 2006). For motifs, the same filtering and refinement procedure was used to make OG-specific models with high specificity and selectivity (Figure 1).

### Inferring repeat dynamics from repeat tree - gene tree reconciliation

We compared repeat trees with gene trees to infer repeat unit duplications and losses, analogous to how gene duplications are inferred from the reconciliation of gene- and species trees. To obtain repeat trees that could be reconciled with the gene tree, we made phylogenetic trees from the repeat units with IQ-TREE (Nguyen et al. 2015) for every OG. Comparison of the repeat tree and gene tree was done by TreeFix (Wu et al. 2013) with additional downstream processing. By using gene trees instead of species trees and one-to-one orthologs, we could include fast-evolving protein families which increased the scope of our analysis: 71% (3515 OGs) of our dataset has gene duplications since the ancestor of vertebrates. Spurious evolutionary events were prevented by allowing for rearrangements in the repeat tree during reconciliation. TreeFix finds a topology of the repeat tree that minimizes events without significantly lowering the tree’s statistical support based on sequence information. In addition, duplications with a consistency score of zero were removed. These two strategies drastically lower the total number of post-ancestral duplications (70.825 to 13.564). As a relative measure of repeat region evolution, the PRD score is calculated for each OG (Supplementary Table 1). PRD score = (x − u) / n; with x = duplications since most recent common ancestor; u = mean duplications in the full dataset, n = number of proteins.

As additional resources, OG-specific repeat unit HMM profiles and (reconciled) repeat trees are available on GitHub, as well as extended data tables about detected repeats and inferred evolutionary events. https://github.com/ivanbelzen/PhyRepID/

### VCAM1 case study as manual verification of Ig repeat detection and reconciliation

As proof of principle, a moderately fast evolving OG (#242) was used as a case study in which we manually checked the repeat annotation and reconciliation process (Supplementary Note 1). Vascular cell adhesion protein 1 (VCAM1) has an immunoglobulin repeat region consisting of six repeat units on average (Figure 2). Comparing repeat annotation using VCAM1’s OG-specific repeat unit HMM with using the best-matching HMM from Pfam shows the severe underdetection by the generalized Pfam model. The repeat annotation after our refinement steps is in accordance with the manually curated UniProt annotation of human (P19320) and mouse orthologs (P29533) (The UniProt Consortium 2019) and the multiple sequence alignment. In addition, we confirmed the placement of duplications on the gene tree with manual reconciliation using repeat unit homology, presence/absence patterns and repeat order within the protein. Note that especially the improvement in annotation greatly reduces differences between proteins and therefore prevents inference of false duplication and loss events, e.g. by filling in gaps, extending partial hits (leading to better repeat trees) and removing spurious (i.e. false) annotations.

**Figure 2:**
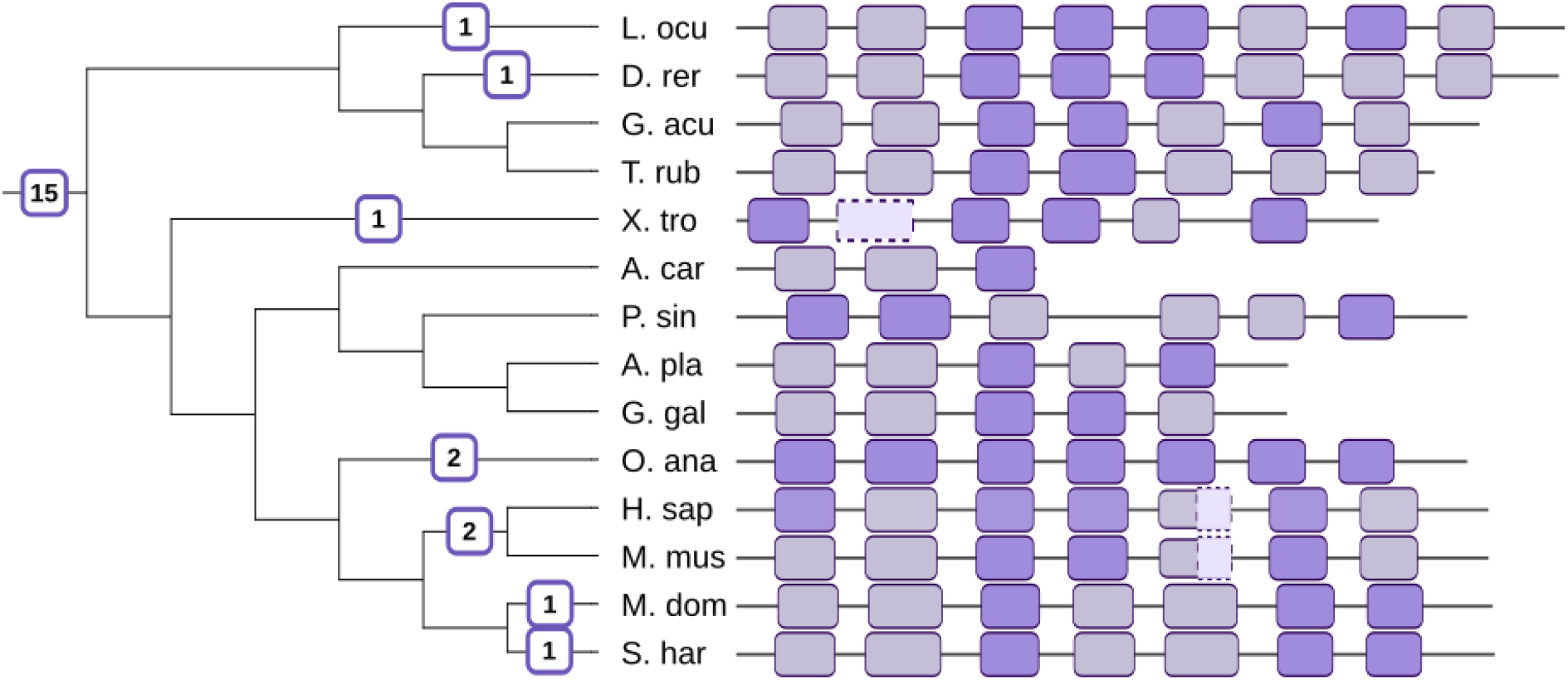
VCAM1 immunoglobulin repeat region annotation before and after refinement. Gene tree with repeat duplications (left) and corresponding orthologs with annotated repeat region (right). Annotation with the best-matching Pfam HMM (ig_3) and applying the default domain model hit cut-off, called the Pfam gathering threshold, results in severe underdetection (dark purple annotated regions are only detected). Also, permissive annotation allowing repeat units below the threshold (light purple annotated regions detected additionally) still missed two partial repeat units in the human/mouse proteins and one repeat unit in the frog protein (annotated regions with dotted lines). VCAM1’s OG-specific repeat unit HMM detected all displayed repeat units, in accordance with manually curated UniProt annotation of human (P19320) and mouse orthologs (P29533). In doing so, differences between proteins are greatly reduced and false inference of duplication and loss events was prevented.

## Results and discussion

### The PRD score: ranking OGs in terms of repeat evolution

From a phylome of repeats we derive the PRD score that reflects the relative speed of repeat evolution and makes it possible to compare vertebrate orthologous groups (OGs) with different sizes (Figure 3). In total, the PhyRepID pipeline contains 4939 OGs with 60.028 repeat proteins of three or more domains/motifs. A ranking of these protein families is made according to their PRD score to compare their relative repeat evolution (Supplementary Table 1).

**Figure 3:**
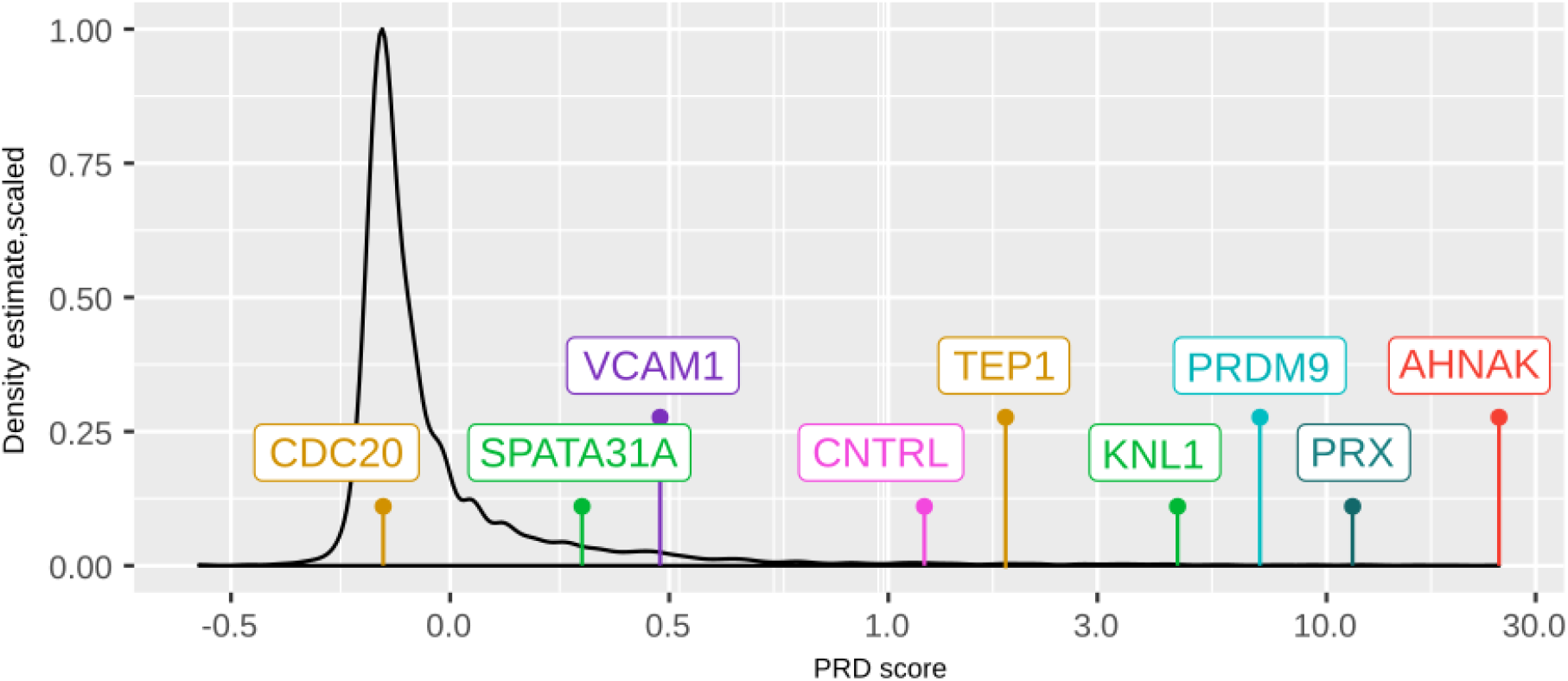
PRD score landscape. Distribution of OGs with PRD scores (x-axis symlog) shows the majority of OGs is conserved (PRD score < 0) with a long tail of intermediate to fast-evolving repeat protein families.

Repeat proteins that are strongly conserved or very dynamic have PRD scores that reflect their previously described behaviour. Negative PRD scores can be interpreted as “more conserved than average”, such as the cell cycle protein CDC20 (−0.15) that has a constrained beta propellor fold (Yu 2007; Xu and Min 2011). As expected, KNL1 (#20, Figure 4a) and PRDM9 (#14) have especially fast evolving repeats (Schwartz et al. 2014; Tromer et al. 2015; Roy et al. 2020). Strikingly, most fast evolving repeat proteins have not been previously reported as such. Novel interesting candidates include #1 AHNAK and #8 PRX which share a neural functional annotation (Masuho et al. 2008; Salim et al. 2009). AHNAK (desmoyokin) is expressed in diverse cell types: from endothelial cells forming the blood-brain-barrier to cardiac muscle cells and myelinating Schwann cells (Salim et al. 2009). AHNAK was described to have a large conserved central repeat region, short N-terminal domain and longer C-terminal domain (Shtivelman et al. 1992). PRX (periaxin, Figure 4b) is also associated with the maintenance of myelin and has a similar central repeat region as AHNAK, but it has a complementary expression in Schwann cells (Salim et al. 2009). Note that the apparent gene prediction problems in the AHNAK and PRX protein families potentially influence the repeat region analysis (Tørresen et al. 2019). For PhyRepID this effect is fortunately small because the PRD score uses inferred duplications and records, but does not incorporate loss events precisely for this reason (see methods).

**Figure 4:**
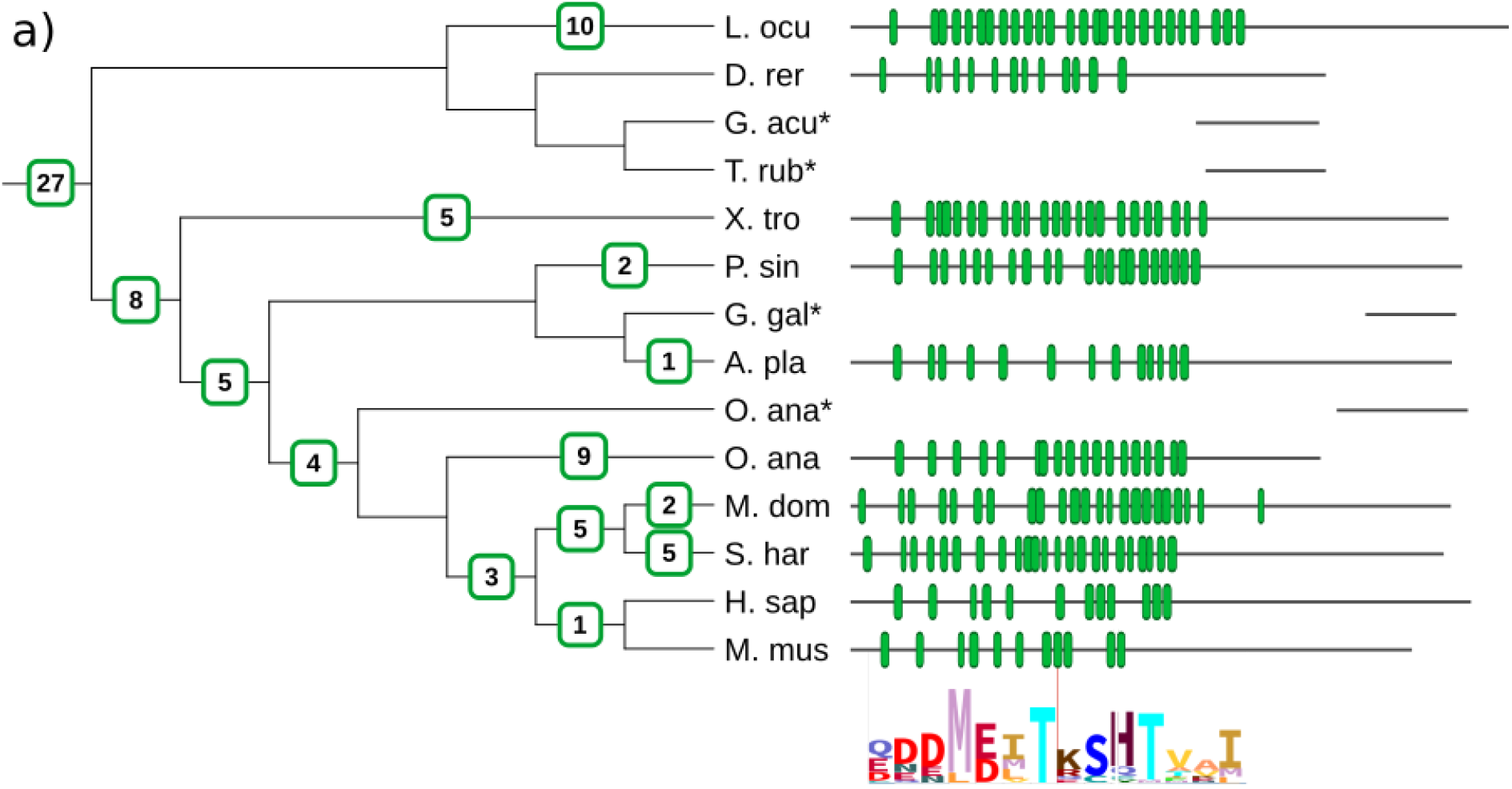
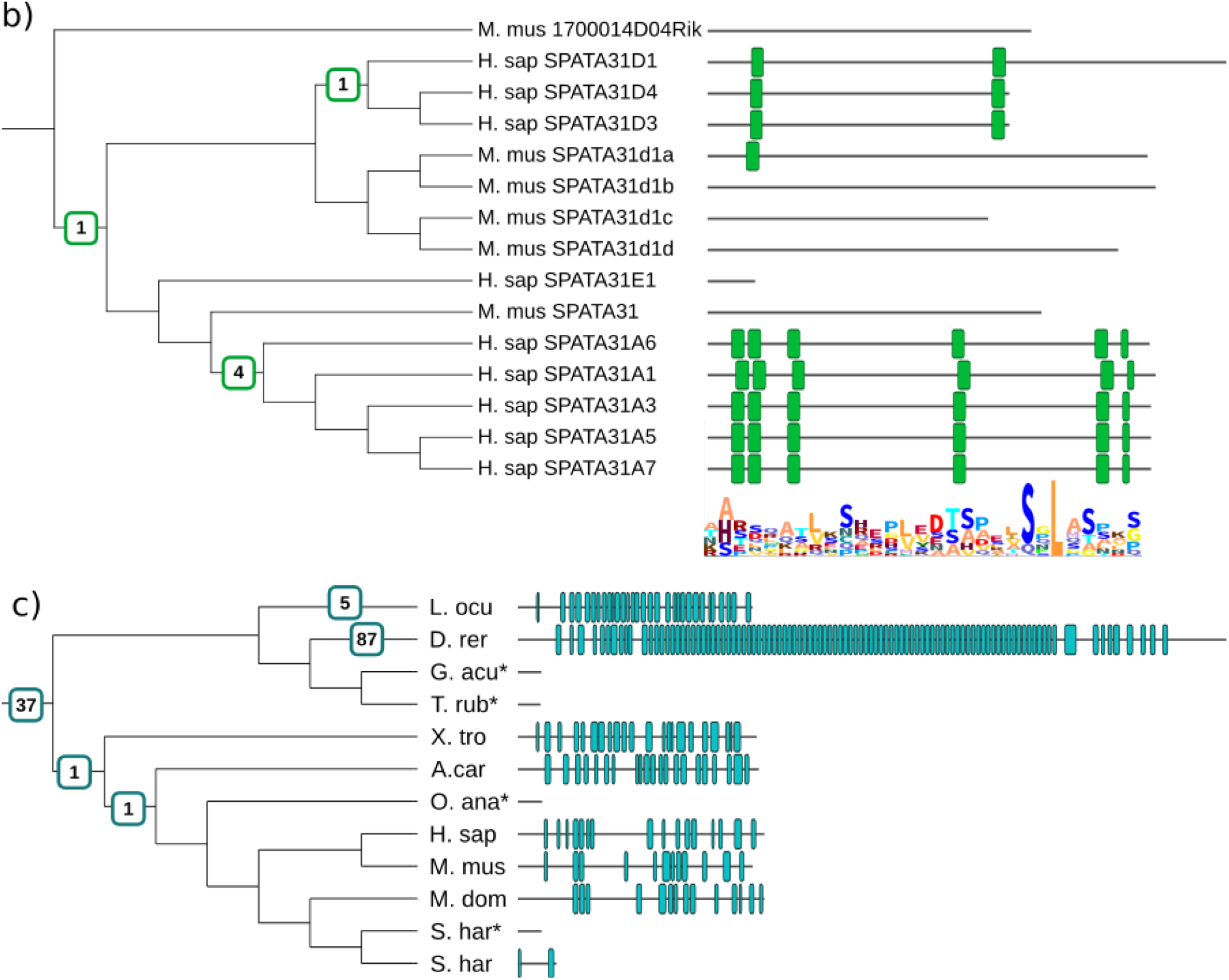
Examples of relevant repeat protein families. KNL1 (a), SPATA (b), PRX (c). Gene trees with repeat duplications (left) and corresponding orthologs with annotated repeat region (right). For KNL1 and SPATA the sequence logo of the motif is shown below/next to the tree/annotated repeats. * denotes gene prediction issue, fragments of N-terminus shown at approximate homologous region

### The SPATA31 protein family shows recent human-specific motif duplications prior to subfunctionalization

To look for recent human repeat evolution, OGs were compared based on duplications only on the human branch exemplifying how the pipeline can be used to study lineage-specific evolutionary dynamics (Supplementary Table 3). The SPATA31 (spermatogenesis associated protein 31, Figure 4c) protein family has a high fraction (0.88) of human-specific duplications and PRD score > 0. The SPATA31 proteins bears an interspersed motif repeat that partially matches binding sites for kinases (Gouw et al. 2018). This repeat region consists of 2-6 units in the nine human paralogs and only a single repeat unit in the mouse ortholog. Using the placement of duplications on the gene tree and the homology pattern in the repeat region, we can speculate that motif duplications may have enabled the recent gene duplications of SPATA31A proteins in the human lineage (~98% sequence identity). After this duplication, the tissue expression of the SPATA31A proteins broadened and the proteins acquired a function in UV damage response (Bekpen et al. 2017).

### Rapidly evolving zinc finger repeats and conserved solenoid-type domains - associations between repeat unit class and evolutionary rate

To look for trends, we analysed the long-tail of fast-evolving repeat protein families (Figure 3) to find characteristics and differences compared to the conserved majority (79%, 3908 OGs with PRD score <=0). Based on protein-level functional annotation, the full set of repeat-containing proteins that we identified has a distinct annotation profile compared to all human proteins that is consistent with previous findings (Björklund et al. 2006) (Supplementary Table 4). Functional overrepresentation trends of dynamic repeat proteins (1031 OGs with PRD score >0) compared to a background of all repeat proteins are less pronounced. There is a slight enrichment of proteins associated with regulation of metabolic processes (GO:0019222) (1.49x), gene expression (GO:0010467) (1.44x) and zinc finger transcription factor (PC00244) (1.78x) function (Supplementary Table 4). For the 100 fastest evolving OGs, functional overrepresentation analysis did not provide further insights since many of these repeat protein families are uncharacterized.

Alternatively, characterization of the OGs based on the detected repeats did reveal associations between the repeat unit class and evolutionary speed as measured by PRD score (Figure 5, Table 1). In the 100 fastest evolving OGs, protein families with C2H2 zinc finger repeats were significantly overrepresented (28%, 9.2x enrichment p<0.01). Many of these high ranking zinc finger proteins are functionally uncharacterized and interesting for follow-up research; they could be part of novel complex gene regulatory networks or involved in the arms race with repression of transposons (Jacobs et al. 2014; Yang et al. 2017). In addition to these very rapidly evolving OGs, zinc finger OGs on average have a higher PRD score than other domains or motifs (Δ0.38, p<0.01) (Figure 5, Table 1). The independent domain folding and beads-on-a-string structure of zinc fingers (or similar nucleic acid binding domains) might contribute to an easier loss or gain of repeat units (Kajava 2012; Schüler and Bornberg-Bauer 2016).

**Figure 5:**
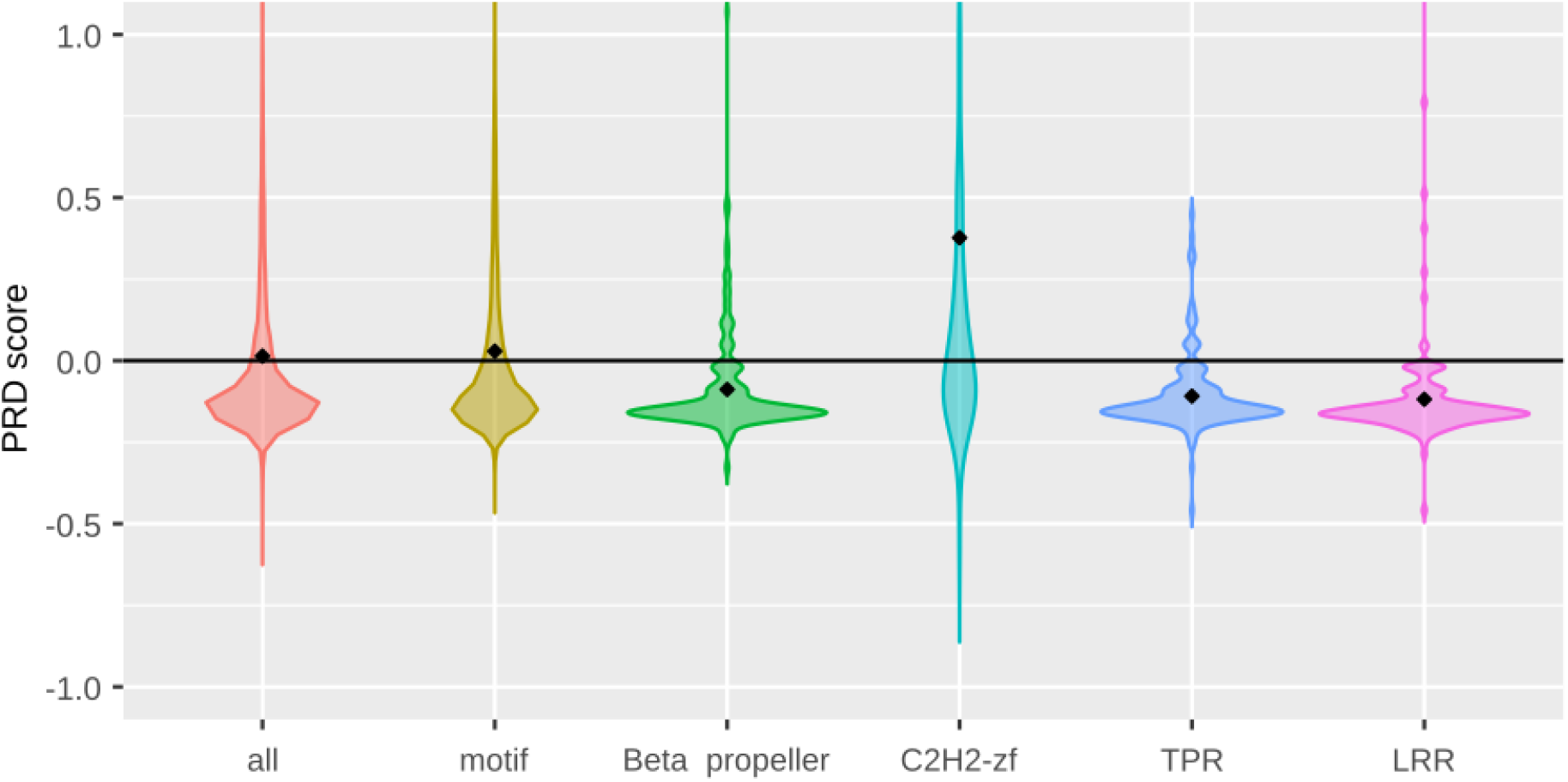
Comparison of PRD score distributions across the most prevalent repeat unit classes. Protein families with C2H2 zinc finger repeats evolve significantly faster compared to the full dataset, whilst beta propellor, TPR and LRR show significantly more conserved evolution. Motif repeats have a similar distribution as the full dataset.

De novo detected motif repeat proteins are neither enriched nor depleted in the 100 fastest evolving OGs compared to the full dataset as they comprise about half in both sets (Table 1). Although the mean PRD score of motifs is slightly higher than domains (Δ0.03, p<0.01), the biological significance of this difference is unclear. There is also no apparent difference in repeat evolution speed between motif repeats and Pfam domain repeats according to their PRD score distributions (Figure 5). Comparing functional annotation of proteins with dynamic motifs shows a significant depletion of proteins associated with cell adhesion (GO:0007155) (0.16) and the extracellular matrix (PC00102) (0.07) compared to a background of all repeat proteins, whilst they share functional overrepresentation trends with dynamic domain repeats. Hence, motif repeat proteins form a heterogeneous superset which potentially contains unrecognized subclasses with different dynamics.

In contrast to these rapidly-evolving examples, proteins with repeats consisting of beta propeller, TPR, LRR or Ank domains have significantly lower mean PRD scores (Figure 5, Table 1). Hence these repeat regions are more conserved than average, as expected based on the biophysical constraints of these domain repeats (Andrade et al. 2001; Kajava 2012). Despite these trends, some generally conserved domains still display dynamic evolution in some proteins such as the telomerase associated protein 1 (TEP1) with beta propeller repeat #52 and centriolin (CNTRL/CEP110) with LRR #71. Interestingly both proteins are associated to processes where rapid repeat evolution is seen more often namely RNA binding (TEP1) and meiosis and spindle formation (CEP110) (Poderycki et al. 2005; Gordon et al. 2006).

### Many rapidly evolving repeat protein families are newly reported candidates

To investigate whether protein families in which we reported repeat evolution have previously been identified, the PRD score was compared to other measures of protein evolution. For comparisons with external datasets that only consist of human proteins (Moretti et al. 2014; Schaper et al. 2014; Karczewski et al. 2017), we defined a “dynamic in human lineage” subset of 1588 OGs that have at least one duplication in the amniote branch up to and including the human branch (Supplementary Table 5).

Comparison to the analysis by Schaper *et al.* (2014) shows that only 3% of the OGs where we detect evolutionary dynamics in the human-lineage were identified previously (Table 2). We identified a number of factors that contribute to the underdetection of these dynamic repeats in previous efforts. First, Pfam domains were used at default cut-off values without heed to inter-domain homologies (i.e. Pfam clans). This leads to underdetection of domains due to insensitivity, as well as domain identity differences between species (see discussion). Second, the extent of the orthology detection by Schaper *et al* was limited as only for 36% of the OGs (1761 of 4939) in our dataset a human-ortholog pair was available in their dataset. Third, whilst Schaper *et al.* approximates phylogeny, they do not utilize it explicitly to study repeat dynamics. Finally, in 92 OGs Schaper *et al*. detects repeat evolution, for 47 OGs of these 92 OGs PhyRepID also detects dynamics in the human lineage. For the 45 other OGs, manual inspection revealed that these repeats are not dynamic. Instead, their bi-species comparisons were inconsistent due to aberrant Pfam domain annotation (Supplementary Note 1). This is also supported by the decrease in variation we observed after domain detection refinement compared to the default Pfam annotation (Supplementary Table 2). For the 1668 proteins where Schaper *et al.* detected conservation, we also find conserved repeat evolution in 82% (1369 OGs with PRD score <0). From this comparison, it seems that PhyRepID detects protein repeat evolution with higher sensitivity and precision than previous large-scale analyses.

**Table 2:** Comparison of our pipeline with Schaper *et al.* (2014) results. Fisher enrichment OR 2.61x (95%CI 1.67-4.08)

| **Table 2: Comparison of our pipeline with Schaper *et al.* (2014) results.** |  |  |
| --- | --- | --- |
| Fisher enrichment OR 2.61x (95%CI 1.67-4.08) |  |  |
|  | **Dynamics in the human lineage** |  |
| Schaper annotation | Yes | No |
| Separated | 47 | 45 |
| Conserved | 476 | 1192 |
| No annotation | 1065 | 2114 |

### Comparison to other measures of rapid evolution reveals independence of the repeat signal

Rapid evolution has been linked to adaptive processes as well as mutational biases. We thus compared proteins with rapidly evolving repeats to other measures of rapid evolution or indications of a change in selection regime (Supplementary Note 2).

We first compared proteins with rapid repeat evolution to a curated set of vertebrate proteins with significantly elevated dN/dS ratios which is strongly suggestive of positive selection. The Selectome database was used to obtain proteins with elevated dN/dS ratios (Moretti et al. 2014). The majority of protein families with rapidly evolving repeats were absent from Selectome. Instead, we observed that repeat proteins that are under positive selection according to Selectome have a slightly lower mean PRD score (−0.028 vs 0.026 p<0.01) and are depleted in the dynamic human lineage set (OR 0.63, 95%CI 0.54-0.74).

We additionally compared known human protein coding variation as this is indicative of relaxed purifying selection. Hereto the ExAC database was used (Karczewski et al. 2017). The sensitivity for missense mutations and loss-of-function mutations is slightly higher for the proteins in the dynamic human lineage set compared to the full dataset (mean missense z-score 1.65 vs 1.26, p<0.01, mean pLI score 0.53 vs 0.46, p<0.01), suggestive of increased purifying selection. Yet, the 100 fastest evolving protein families do show somewhat relaxed constraints (mean missense z-score 0.62 and pLI 0.30, p<0.01).

Third, evolutionary dynamics in repeat proteins could be related to gene duplication which is a known driver of adaptive evolution (Kondrashov 2012). We found that significantly more gene duplications were present in the 100 OGs with fastest repeat evolution (mean 3.29 vs 2.40). In addition, protein families with rapidly evolving repeats are more likely to have complex orthologies (40% vs 9.8%, OR 6.6x, 95%CI 4.23-10.1, p<0.01). Complex orthology cases are protein families without a bony vertebrates common ancestor node in the gene trees from ENSEMBL Compara, despite having domains indicative of older evolution ancestry. This means that even the advanced Compara pipeline is unable to solve their precise relations due to promiscuous nested duplications.

Fourth, to study the involvement of genome organisation we compared the location of human orthologs with segmental duplication (Duplicon) regions (835) (Marques-Bonet and Eichler 2009). There is a slight enrichment in the dynamic human lineage set for OGs with human orthologs located in duplicon regions (299 of 1588, OR 1.22x 95%CI 1.03-1.43 p<0.05) and also 20 of the 100 fastest evolving OGs (OR 1.23x, n.s.) are located here. Whilst gene duplication and genome organisation factors might influence the speed of protein repeat evolution, they seem in itself unable to provide an explanation for all cases of rapid repeat evolution so there are likely other factors involved.

## Discussion

The PhyRepID pipeline enables large-scale, quantitative comparison of the evolutionary dynamics of repeat proteins across vertebrate species (Figure 1), assessing 4939 groups of orthologous repeat proteins (OGs). We identified novel candidates (e.g. AHNAK, PRX) with rapidly evolving repeats by using the PRD score to rank protein families according to repeat evolution. In addition, we are the first to quantify differences between known conserved and rapidly evolving OGs (i.e. CDC20 versus KNL1 and PRDM9). We find that protein repeat evolutionary dynamics is influenced by and/or characteristic for the repeat unit type. As expected, protein domain repeats known for cooperative folding are mostly conserved (i.e. beta propellor, TPR, LRR, Ank) (Kajava 2012). Strikingly, C2H2 zinc-finger proteins show rapid evolution and are significantly over-represented in the 100 fastest evolving OGs. Motif repeats form a highly heterogeneous group with both rapidly evolving and conserved protein families showing a similar PRD score distribution as domain repeats. This suggests that motif sequences of intermediate length (10-30 aa) that are currently uncharacterized might have interesting evolutionary properties that warrant further research to aid in their classification.

### Progress and limits in characterization of repeat evolution

The PhyRepID pipeline quantifies repeat evolution using comparative phylogenomics of orthologous groups (OGs). Our three main contributions are (1) detection of a wide scope of repeats consisting of either Pfam structural domains or *de novo* detected linear motifs, (2) achieving high sensitivity and precision of detection through optimization of repeat unit Hidden Markov Models (HMMs) for each OG, (3) using a novel two-step phylogenetic tree reconciliation approach by first comparing repeat trees to gene trees prior to inferring repeat duplications on the species tree.

To assess evolutionary events in the repeat region as accurately as possible, we found that precise domain annotation is of high importance. First, including *de novo* motifs strongly increased the number of OGs in which repeat regions could be detected: 2676 OGs would not have been included if only Pfam domains were considered. Second, using OG-specific repeat unit HMMs prevents false duplications and losses by filling in gaps from undetected or partial repeat units, as well as removing spurious annotations. Hence, although Pfam is a valuable resource, relying on generalized domain HMMs has caveats for comparative genomics. Protein domains have varying degrees of sequence divergence within and between protein families (Andrade et al. 2001; Stirnimann et al. 2010) which is problematic for accurate detection with generalized Pfam HMMs and can lead to missing protein domain occurrences. In an effort to circumvent this, multiple similar HMMs were created for homologous domains and subsequently grouped in Pfam clans (Punta et al. 2012). Also, for some domains that are often tandemly repeated (e.g. TPR, LRR, Ank) Pfam contains HMMs which are multi-models of 2-3 repeating units, because longer HMM profiles can improve detection sensitivity (El-Gebali et al. 2019). However, multiple homologous HMMs does not resolve issues with sequence divergence and complicates uniform repeat detection within a protein family, as observed by a variety of best-matching domain HMMs in orthologs (Supplementary Note 1). Also, multi-models are problematic for assessing the number of repeat units and therefore quantification of repeat dynamics. In our approach, we group the best-matching HMMs on clan level and build an OG-specific HMM matching one repeating unit of the homologous repeat region in the protein family. Making these optimized OG-specific repeat unit HMMs improved the sensitivity and precision of detection, and prevented the inference of spurious duplications and losses for 1 in 10 Pfam OGs (Supplementary Table 2).

Using a two-step phylogenetic tree reconciliation to analyze repeat evolution enables us to distinguish between evolution on the gene level and repeat level. Incorporating information from gene trees prevents the inference of spurious evolutionary dynamics. As a consequence, repeat evolution can be studied per branch of the species tree, as shown by the identification of recent duplications on the human branch of the SPATA31 protein family (Figure 3). In addition, the use of gene trees instead of one-to-one orthologs allows for the inclusion of rapidly evolving OGs with recent gene duplications since the ancestor of vertebrates. This is especially relevant as it turns out there is an evolutionary and presumably functional relation between gene duplication and protein repeat dynamics (largely driven by zinc fingers).

However, in order to reconcile the repeat tree a correct gene tree is of importance. We retrieved gene trees from ENSEMBL Compara (Zerbino et al. 2018) and noticed that ~10% of OGs have a truncated gene tree without a node in bony vertebrates. These truncated gene trees are indicative of a failure to identify older orthologous relationships by the Compara pipeline due to spuriously dividing larger orthologous groups into smaller ones. In the most extreme cases, human genes that stem from the vertebrate ancestor were not present in our dataset, because the pipeline requires at least one human-mouse ortholog. As a result, some rapidly evolving protein families with older orthologs undefined by ENSEMBL were not included in our analysis, e.g. the KRAB-zinc finger family ZNF91, ZNF93, ZFP57, ZFP568 which is associated with transposon repression (Jacobs et al. 2014; Yang et al. 2017). Note that missing ancient orthologs also leads to an underestimation of repeat evolution in highly dynamic protein families and possibly obfuscates the relationship between gene - and repeat duplication. Despite the underdetection that gene tree truncation leads to, we find that truncated OGs are over-represented in the 100 fastest evolving OGs (6.5x p<0.01) which also suggests a possible relationship between gene duplication and repeat dynamics. Possibly, gene tree prediction can be improved by incorporating information from the repeat trees, i.e. by co-estimation of the repeat and gene tree or by deriving a consensus or ancestral repeat using concatenated alignment.

### Comparison of rapid repeat evolution to other rapid evolutionary processes

We have found that fast protein repeat evolution is a source of evolutionary variation not measured by established methods for detecting rapid protein evolution. Although there is some relation to some of these measures (especially gene duplication), there are also conspicuous absent relations such as the lack of significant overlap with positive selection as inferred from high dN/dS ratios. Possibly this is because DNA-based methods are limited to short evolutionary timescales (e.g. human-primate lineage) (van der Lee et al. 2017) and require highly similar sequence (Persi et al. 2016; Schüler and Bornberg-Bauer 2016) considerable sequence divergence. By using phylogenomics, the PhyRepID pipeline can identify protein repeat evolution across longer timescales than is possible with DNA-based measuresless of evolution like dN/dS. Note that the PRD score does not have a null expectation and therefore we cannot statistically separate neutral from adaptive evolution, however we are able to observe the absence of (strong) purifying selection. Since the PRD score is formulated relative to the average duplications of the dataset, it is a conservative estimate specifically aimed at detecting fast-evolving protein families.

It is tempting to speculate that repeat regions can provide proteins with evolutionary degrees of freedom, since the repeat region as a whole is subject to functional constraints rather than the individual units (Andrade et al. 2001) due to their homologous sequences (Björklund et al. 2006). For example, in KNL1 the rapid evolution of the motif repeat region seems to be adaptive and result from balancing the strength of motif binding and responsiveness leading to a varying selection pressure (Roy et al. 2020). In addition, because (i) there is also no clear mutational bias, (ii) most repeats are stably conserved, and (iii) purifying selection seems to be active on proteins with dynamic repeats (as inferred from ExAC and Selectome), it is possible that those repeats that are rapidly evolving, do so for adaptive reasons which still would need to be elucidated.

### Conclusion and outlook

The PhyRepID pipeline and derived PRD score enable comparison of repeat proteins in terms of their evolutionary dynamics since the common ancestor of vertebrates. Especially the high-ranking uncharacterized proteins are interesting candidates for more in-depth analysis of their biological function. The extent of the dataset resulting from this pipeline also makes it possible to study protein repeat evolution in much more detail. Information on evolutionary events is available as gene tree annotation and could be used to study branch-specific evolution or associations between the timing of gene and repeat duplication. This can aid in generating hypotheses about evolutionary pressures and possibly adaptive processes in protein repeats, as well as in resolving existing biological questions about (genome) evolution and speciation. The PhyRepID pipeline can also be applied to study repeat evolution in other groups of species, e.g. the Drosophila genus. Applying these to other taxa would join our already extensive list of candidate genes to elucidate biological causes for these repeat dynamics.

## Materials and methods

The PhyRepID pipeline is available as Snakemake workflow on GitHub https://github.com/ivanbelzen/PhyRepID together with Supplementary figures, tables and extensive documentation. The repository contains the scripts for data-processing, figures and statistical analyses, as well as data dumps consisting of OG-specific repeat unit HMMs and (reconciled) repeat trees.

### Data collection: construction of a dataset with orthologous groups

Orthologous proteins were acquired from ENSEMBL Compara v91 (Zerbino et al. 2018) using a SPARQL query on the EBI endpoint on 12-04-2018. All homologs were retrieved from *Homo sapiens* (human) protein coding genes with each of the 13 other vertebrate species: *Mus musculus* (mouse), *Monodelphis domestica* (opossum), *Sarcophilus harrisii* (Tasmanian devil), *Ornithorhynchus anatinus* (platypus), *Anas platyrhynchos* (duck), *Gallus gallus* (chicken), *Gasterosteus aculeatus* (stickleback), *Takifugu rubripes* (takifugu), *Pelodiscus sinensis* (turtle), *Anolis carolinensis* (anole lizard), *Xenopus tropicalis* (frog), *Danio rerio* (zebrafish), *Lepisosteus oculatus* (spotted gar).

The resulting set of orthologous groups (OGs) were filtered. The human protein coding gene should have:

- At least one ortholog in mouse
- Orthologs for both or no species in a pair: human-mouse, opossum-tasmanian devil, duck-chicken, stickleback-takifugu. These species pairs are made based on divergence times of approximately 80 million years (Hedges et al. 2015). This filtering should prevent gene prediction errors to cause duplications and losses, and give more confidence to the repeat tree because there are two proteins with higher sequence identity.
- A maximum of three orthologs in a single species, which will be paralogous to each other. Note that the number of human paralogs in an OG is not restricted. This filtering was done for computational reasons.

This resulted in a dataset of 11.755 OGs of which gene trees and protein sequences were retrieved via the ENSEMBL API. The smallest gene tree containing all orthologs was extracted. In subsequent steps, the OGs are identified by a concatenation of the gene tree ID and their human ortholog ENSEMBL gene ID.

### Repeat detection

#### Pfam domain repeat detection

A liberal HMMscan was performed for all OGs against Pfam-A 31.0 (Finn et al. 2016) with HMMER3.1.b2 (Finn et al. 2011) using a minimum sequence bit score of 12.5, which is lower than all sequence gathering thresholds, and 0 as domain threshold (*-T 12.5 --domT 0*). Filtering is done subsequently using the models’ sequence gathering thresholds, which are manually curated bit score cutoffs for all hits within a sequence (Punta et al. 2012). The best-matching Pfam HMM for each Pfam clan is selected based on the highest sequence bit score over all sequences in the OG. A permissive definition for repeat proteins is used: a minimum of three repeat units in a human protein and at least one in a mouse protein, resulting in 2263 OGs with a Pfam repeat region.

After the initial repeat detection, OG-specific repeat unit HMMs were made to improve the detection sensitivity and specificity. Repeat units detected by the best-matching Pfam HMM were extracted using target coordinates without padding, after which a multiple sequence alignment (MSA) was made using MAFFT L-INS-I *(--localpair --maxiterate 1000)* which uses pairwise alignment (Katoh et al. 2005). Profile HMMs were constructed from the MSA using default settings *(--fast --symfrac 0.6)* (Eddy 1998). Next, an iterative process ran until convergence for each of the OGs, consisting of the following steps: (1) HMMscan was performed with the profile model, (2) an MSA was made of the hits and from this MSA, (3) a new profile was constructed. This process was repeated until convergence as indicated by subsequent iterations detecting the same repeat units, or a maximum of 20 cycles was reached. To prevent profile drift to short lengths for highly repetitive models, the iteration was also terminated if the profile reached a length of <20 amino acids. After termination, a final MSA was made with envelope coordinates for the domain sequences and a padding of +− 5 amino acids. This slightly enlarged the repeat sequence and provided more information for constructing the repeat tree.

#### Motif repeat detection

The MEME package was used for *de novo* detection of motifs in OGs in which no Pfam domain repeats were found (Bailey and Elkan 1994). A search was performed for the occurrence of any number of ungapped sequences (*-protein -mod anr*) of a length of 10 to 30 amino acids (*-minw 12 -maxw 30*) that is present a minimum of 4 times in the OG (*-minsites 4*). The search is terminated after one such motif is found, and otherwise if 300 cpu seconds or 10 iterations have passed (*-time 300 -maxiter 10*). As input, a fasta file with the protein sequences of an OG are provided (*-protein -maxsize 10000000*) with regions matching Pfam domains removed from the sequence to prevent matching these during motif detection. Note that this could not be solved by masking, since it lead to the detection of motifs consisting of X’es.

Filtering was subsequently done based on the same criteria as used for Pfam domains: requiring a minimum of three repeat units in a human protein and at least one in a mouse protein resulting in 2676 OG-motif combinations. After motif detection, the OGs also underwent the same refinement procedure as in the Pfam repeat detection, during which OG-specific repeat unit HMMs are made.

### Tree reconciliation: inferring duplications and losses

Evolutionary events (i.e. duplications and losses) were inferred for each OG from both the Pfam domain and motif datasets via phylogenetic tree reconciliation of the repeat tree with the gene tree. Gene trees for each OG were retrieved from ENSEMBL and parsed as previously described. For the repeat tree, a maximum likelihood (ML) tree was constructed from the repeat sequences in the OG. A species tree for the 14 vertebrate species was manually constructed based on the ENSEMBL species tree (Zerbino et al. 2018). Next, the species tree was manually annotated with taxonomic identifiers for the nodes and divergence times as branch lengths with the help of TimeTree.org (Hedges et al. 2015).

#### Repeat tree construction

IQ-TREE is used to construct ML phylogenetic trees from MSA of the repeats (Nguyen et al. 2015). Using the recommended optional settings: UBoot ultrafast bootstrapping (Hoang et al. 2018) with 1000 replicates and ModelFinder (Kalyaanamoorthy et al. 2017). IQ-TREE handles duplicate sequences by removing them during the analysis and adding them later with a bootstrap value of 0. The unrooted ML tree was used for further analyses and is subsequently referred to as the ‘repeat tree’.

TreeFix was used to reconcile repeat trees with gene trees (Wu et al. 2013). This algorithm takes the MSA into account while minimizing a cost function (both duplications and loss have a cost of 1.0). This is accomplished by navigation through a ML landscape to find a reconciled tree, subject to the constraint that it is as likely as the ML tree provided. The output from TreeFix was algorithmically adjusted to remove spurious duplications at terminal nodes with a custom python script using the ETE package that allowed for rearranging duplications with a consistency score of 0. A consistency score of zero means that there is no overlap in proteins in the left and right branches after the duplication, hence the chance is very small that this is a valid duplication event

#### Inference of evolutionary events: duplications and loss

The ETE toolkit was used to count evolutionary events by performing strict reconciliation of repeat trees with gene trees in which rearrangements are not allowed (Huerta-Cepas et al. 2016). As output, gene trees are annotated with the number of duplications on each node, and a list is returned with the duplications per node of the species tree together with a taxonomic identifier.

Additionally, gene duplications and losses are inferred by reconciling gene trees with the species tree. Combining the two reconciliation steps makes it possible to map evolutionary events from the repeat tree to the species tree by matching their taxonomic identifier. As a result, duplications and losses can be compared between OGs.

### Quantification of evolutionary dynamics: the duplication score

Our main research goal is to come to an integrative score of how fast a repeat region evolves. Hereto only duplications are considered because inference of duplications appears to be more robust than inference of losses. This is due to the common problem in tree reconciliation algorithms of pushing duplications towards the root of the tree (Hahn 2007). These duplications which are placed too early lead to falsely inferring losses, but it does not affect the total number of duplications since this is also driven by the number of copies.

Duplications placed more recent than the root of the species tree are regarded as a signal for evolutionary dynamics since the divergence of vertebrates, they are referred to as ‘net duplications’. The ancestral duplications, which are placed on the root of the gene tree, are subtracted from the total number of duplications. The root of the gene tree is used for this distinction between ancestral and net duplications to prevent giving an artificially low score to OGs with truncated gene trees that do not have an euteleostomi node (see discussion).

The PRD score provides a measure of how fast an OG evolves compared to “the average evolution speed" of OGs in the dataset. The PRD score of an OG is calculated by subtracting the arithmetic average number of net duplications in the dataset from the number of net duplications for that OG, and dividing by the number of proteins in an OG (Supplementary Table 1).

PRD score = (x - u) / n

- x = duplications since the most recent common ancestor
- u = mean duplications in the full dataset
- n = number of proteins in OG

### Downstream analyses

Repeat detection analysis statistics were collected with an inhouse script parsing the raw HMM output (Extended data). As a measure of improvement in the consistency of repeat unit detection, the coefficient of variation in the repeat unit count was used (Supplementary Table 2). The coefficient of variation was compared to the number of repeat units using default Pfam domain annotation, and OG-specific HMMs. A decrease in the coefficient of variation denotes improved consistency. Note that the coefficient of variation is 0 when all orthologs of an OG have the same number of repeat units.

Statistical analyses were conducted using R, of which the output is available as Supplementary Note 2. The Fisher exact test was used to test overrepresentation of certain subsets of OGs compared to the full dataset, for example whether C2H2-zf repeat-containing OGs were overrepresented in the 100 fastest evolving OGs compared to the full dataset. For comparing distributions of continuous variables such as the PRD score of different subsets of OGs, the Wilcoxon test was used. Both the script and output are available on GitHub.

### Human lineage-specific evolutionary dynamics

The PRD score makes it possible to assess relative repeat region evolution of an OG in context of the full dataset of repeat proteins. In addition, it is possible to analyse lineage-specific evolutionary dynamics using the information of where duplications are placed on the species tree.

To study human lineage specific repeat evolution, only the duplications from the amniote node up to and including the human branches were considered (taxon IDs 9606, 314146, 32525, 40674, 32524, 8287, 9347, 1437010, 314147, 32523). For comparison with external datasets that only consist of human proteins, at least one duplication in this amniote-human lineage trace was required. (Supplementary Table 5)

For the human-only analysis, only recent duplications in the human branch (taxon id 9606, protein identifier ‘ENSP*) were considered. The fraction of human only duplications vs the netto duplication was used to rank the OGs on how fast evolving they are in recent human evolution (Supplementary Table 4).

### Comparison with external datasets

From supplementary material of Schaper et al. (2014) we retrieved annotations of bi-species phylogenies from both Pfam and HHRepID (Schaper et al. 2014). In these files pairs of a human protein with its vertebrate ortholog are annotated as strongly conserved, perfectly conserved, strongly separated, perfectly separated or not assigned. To compare Schaper *et al.*’s pairwise annotation to the OGs in our dataset, the pairwise data was projected on OGs based on the proteins in the OG. An OG was annotated ‘positive according to Schaper’ if at least one human protein was regarded as perfectly or strongly separated from an orthologous protein that is also present in our OG.

Selectome is a database of positive selection measured with dN/dS (Moretti et al. 2014). After personal inquiry, a list of human genes under positive selection since the divergence of vertebrates was acquired from Sebastien Moretti on 04-09-2018 (Moretti et al. 2014). For all human proteins in the final dataset, the gene ID was retrieved from ENSEMBL BioMart on 30-09-2018.

The ExAC database v0.3.1 was used to compare our findings to known human variation (Karczewski et al. 2017). It gives scores to proteins that express the level of constraint of synonymous and non-synonymous (missense) mutations by comparison of their expected and observed frequency. In addition, they also score intolerance to heterozygous loss of function mutations, such as premature stop codons, and associated these with disease. The z-scores for synonymous, non-synonymous and loss-of-function mutations were acquired for each human gene, as well as the probability of being intolerant for loss-of-function mutations.

Over-representation tests were done with PANTHER v14.1 using Fisher’s exact test and Bonferroni correction with FDR < 0.05 (Mi et al. 2019). GO-Slim Biological process was used to find a functional enrichment in the full dataset of repeat proteins, searching by ENSEMBL gene id with the human proteome as background. In addition, the same analysis was conducted for the dynamic set using the dataset of repeat proteins as background. (Supplementary Table 3)

Segmental duplication coordinates were retrieved from http://humanparalogy.gs.washington.edu/build37/build37.htm on 29-10-2019. To determine whether human protein representatives of the OGs fall within the segmental duplication areas, this list was intersected with ENSEMBL gene coordinates using *bedtools intersect (Quinlan 2014)*.

### Visualisation of gene trees and protein repeat regions

For visualisation and manual analyses, the ETE toolkit was used to export annotated NHX files. These files were uploaded to the ITOL website (Letunic and Bork 2011), which provides an interactive interface for displaying phylogenetic trees and visualisation of metadata such as duplication counts. Datasets were uploaded to the ITOL websites in the form of generated label and domain templates in order to visualise protein repeat regions next to a gene tree.

