## Supplementary material for "PhyRepID: a comparative phylogenomics approach for large-scale quantification of protein repeat evolution": Supplementary_Note_1_VCAM1_case_study.pdf

**Content**

1. Improving detection with OG-specific repeat unit models prevents inference of spurious evolutionary dynamics
2. Inconsistency in repeat dynamics due to differences in repeat annotation; a comparison of PhyRepID and Schaper *et al.* (2014).
3. A manual example of repeat tree - gene tree reconciliation

Vascular cell adhesion protein 1 (VCAM1) has an immunoglobulin repeat region consisting of six repeat units on average, and one-to-one orthologs are available for all 14 selected species. The VCAM1 orthologous group (OG) shows moderately fast repeat evolution according to its ranking by PRD score (#242 of 4939) which makes it a suitable protein family for a case study. OGs with less dynamics have little to investigate, but rapidly evolving OGs have repeat trees that are difficult to interpret manually.

**1. Improving detection with OG-specific repeat unit models prevents inference of spurious evolutionary dynamics**

In order to achieve high precision and sensitivity of repeat unit detection, Hidden Markov Models (HMMs) were made for each OG to optimally detect its repeat units. This prevents false negatives and partial hits due to the sequence divergence that repeats often show compared to the Pfam consensus (Punta et al. 2012). Comparison of the general Pfam annotation with these OG-specific HMMs shows more consistency between orthologs regarding repeat region annotation. This is reflected in a decrease in the coefficient of variation and in the number of evolutionary events necessary to explain differences in repeat regions (Supplementary Table 2). Hence, false-positive duplications/losses are prevented by filling in gaps due to undetected repeats, extending partial hits and removing spurious annotations.

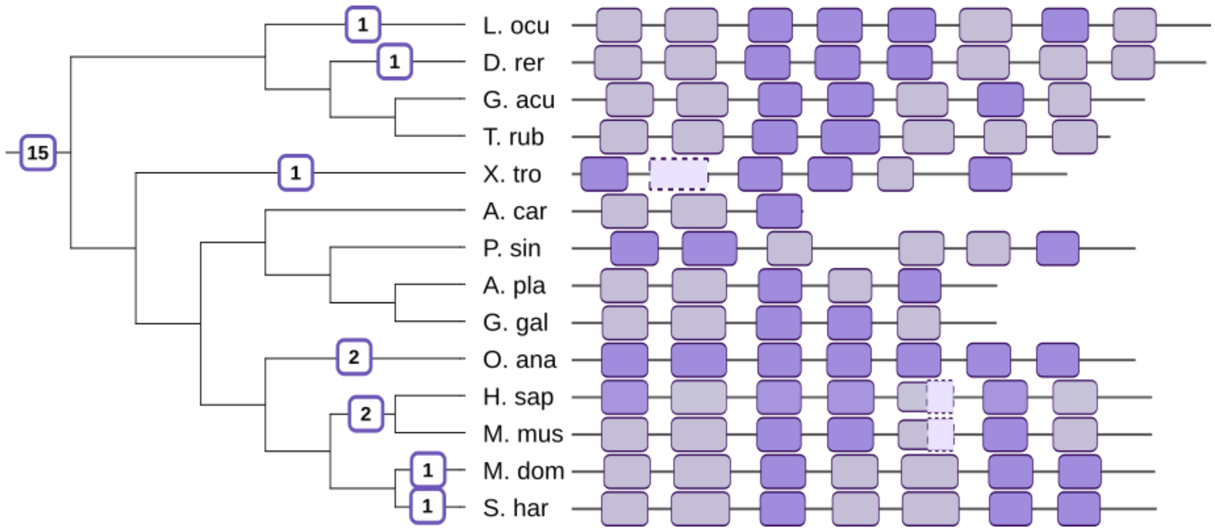


**Figure S1:** *VCAM1 immunoglobulin repeat region annotation before and after refinement. (Corresponds to Figure 2 from main text). Gene tree with repeat duplications (left) and corresponding orthologs with annotated repeat region (right).*

To show the effect of using OG-specific repeat unit HMMs for detection, we compared VCAM1's immunoglobulin repeat region annotation in the final dataset to the initial annotation with general Pfam models (Figure S1). Default annotation using Pfam with the best-matching HMM (ig_3) and recommended domain model hit cut-off (gathering threshold) resulted in severe underdetection (only dark purple regions). When instead permissive settings were used, repeat units below the threshold were found (light purple regions) but this still implied two partial repeat units and gaps (regions with dotted lines). After the OG-specific refinement, the gap in the frog ortholog and the partial hits in human and mouse orthologs were found (Figure S1). The presence of these repeat units was confirmed by manually curated UniProt annotation of human (P19320) and mouse orthologs (P29533), and multiple sequence alignment for the frog repeat unit (Figure S2). However, the gap in the turtle ortholog could not be filled in, hence this gap is possibly a location without Ig domain or highly diverged repeat unit. The phylogenetic tree of repeat units and patterns in clustering also confirm the accuracy of repeat detection. In addition, we confirmed the placement of duplications on the gene tree with manual reconciliation using repeat unit homology, presence/absence patterns and repeat order within the protein (Figure S4, addressed in the next section)

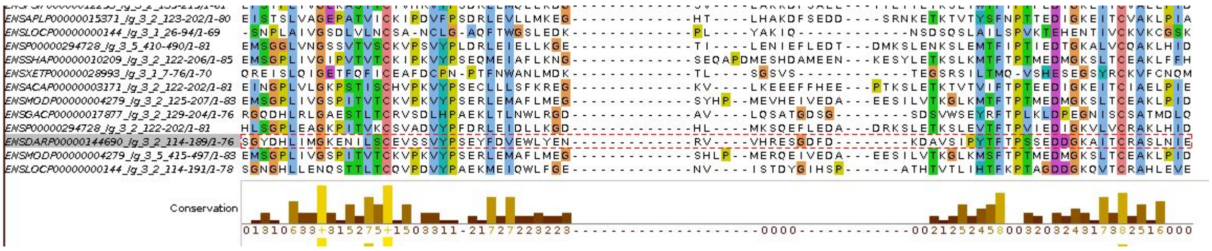


**Figure S2:** *Multiple sequence alignment of VCAM1 repeat units in the 2nd lead order. The repeat unit that was missed in the frog ortholog (highlighted in grey) by default Pfam annotation using the ig_3 model was detected with a OG-specific repeat unit HMM tailored to VCAM*1.

**2. Inconsistency in reported repeat evolution due to differences in repeat annotation; a comparison of PhyRepID and Schaper *et al.* (2014)**

Comparison of repeat evolution as inferred by PhyRepID and Schaper *et al.* (2014) highlighted the improved sensitivity of our approach, as only 3% of the OGs with dynamics in the human lineage were reported previously (Schaper et al. 2014). Only 1761 OGs were present in both datasets and in 92 OGs Schaper *et al.* detects repeat evolution. PhyRepID also detects dynamics in the human lineage for 47 OGs, however for the remaining 45 OGs we find that some are likely false positives due to aberrant Pfam domain annotation.

Although VCAM1 is positive in both datasets, the evolutionary dynamics identified by both approaches are different. The phylogenetic tree reconciliation approach used in PhyRepID is explained in the next section. Schaper *et al.* uses bispecies comparison of repeat region annotations from a human protein and an ortholog from another species to infer ‘separation of the repeat regions’. This approach is highly dependent on the exact matches of Pfam domains and therefore confounded by aberrant Pfam annotation. Unfortunately details are lacking regarding the exact repeat annotation used; only the protein identifiers, Pfam model and conclusion. For the VCAM1 species pairs in which either conservation or separation is detected, Figure S3 displays the annotation using the given Pfam models and the default detection thresholds. Schaper *et al.* concludes that the V-set domain is conserved between human and mouse; and separated for human-frog and human-turtle (Figure S3, top). In addition, separation was detected for human-platypus using the ig model (Figure S3, bottom). Note that multiple Pfam models are used to compare the human VCAM1 protein with its orthologs. This is probably due to multiple Pfam domains hitting the same location (Figure S4). These Pfam models are often homologous but differ in what repeat units they match best within a protein and between species. Manually curated Pfam clans were made to counter this issue by grouping homologous models based on similarities in sequence, structure and function (Punta et al. 2012). Our approach used in PhyRepID prevents using multiple models for comparison by selecting the best-matching model of each clan for the OG followed by the refinement steps to make OG-specific repeat unit HMMs. As a result, the repeat region of VCAM1 is more consistently annotated (Figure S1) which prevents inference of spurious evolutionary dynamics.

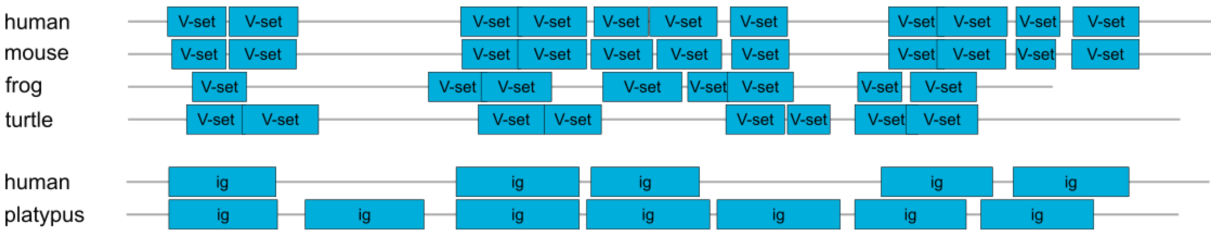


**Figure S3:** *Annotation of VCAM1 proteins and Pfam models which were indicated to be separated or conserved in the analysis by Schaper et al. (2014).*

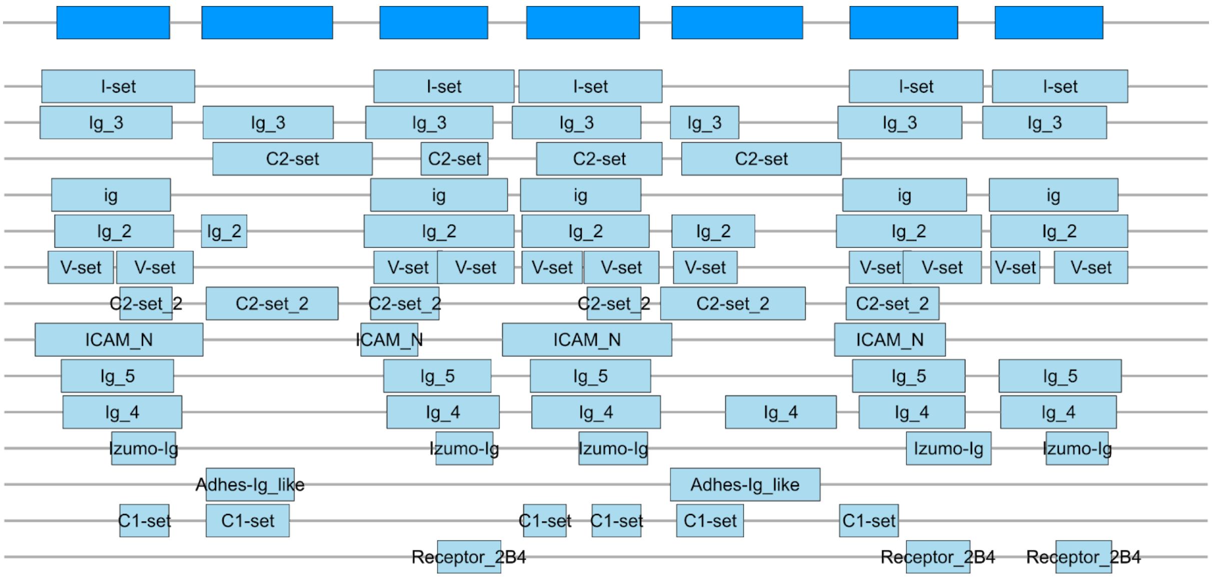


**Figure S4:** *Annotation of the human VCAM1 repeat region with its significant Pfam model hits (light blue) and the repeat region as defined in our dataset at the top (darker blue).*

### 3. A manual example of repeat tree - gene tree reconciliation

The two-step reconciliation used in PhyRepID infers evolutionary events in the repeat region by reconciliation of repeat trees to gene trees, and gene trees to the species tree. Analogous to gene trees, repeat trees are hypotheses of how repeat units are related to each other. In case of a perfectly conserved repeat region, all species's first, second, third etc. repeat unit would cluster together.

For this manual reconciliation example of VCAM1, we identified clusters of homologous repeat units in the tree using patterns in the presence of the majority of species, the lead order in which they occur in their repeat region (analogous to synteny) and bootstrap values (Figure S5). Note: clustering is not used in the pipeline but only for illustrative purposes.

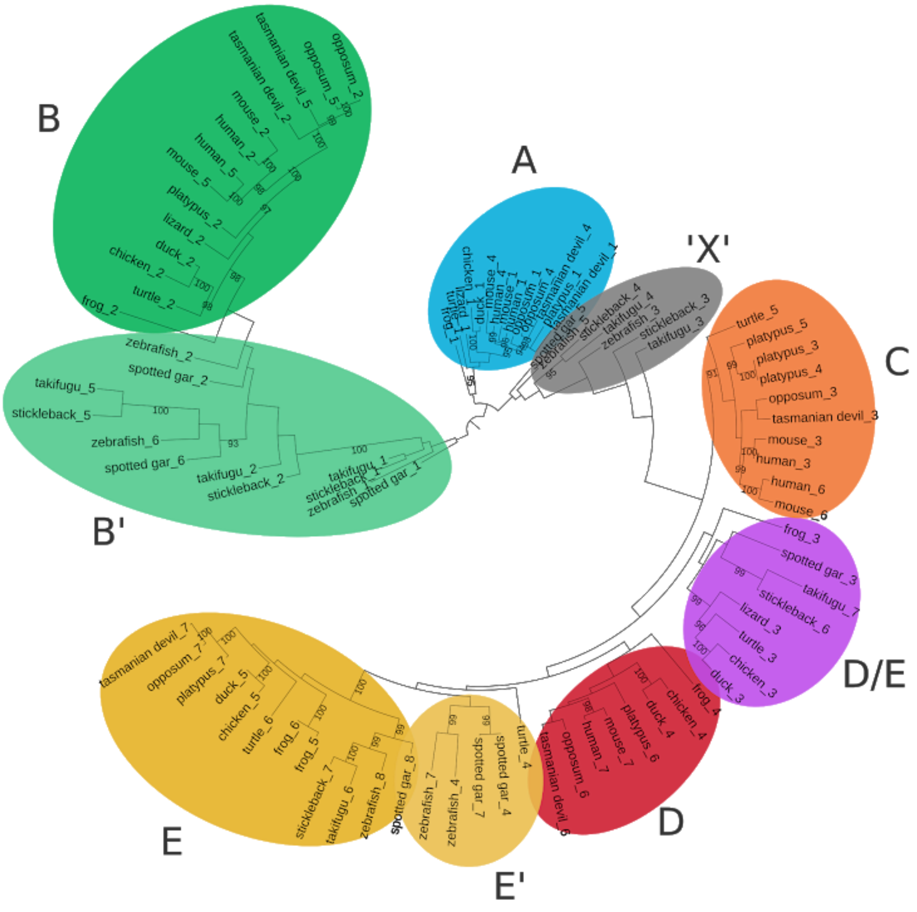


**Figure S5:** *VCAM1 repeat tree with coloured and annotated clusters of repeat units based on bootstrap values, presence/absence patterns in orthologs, lead order and manual inspection.*

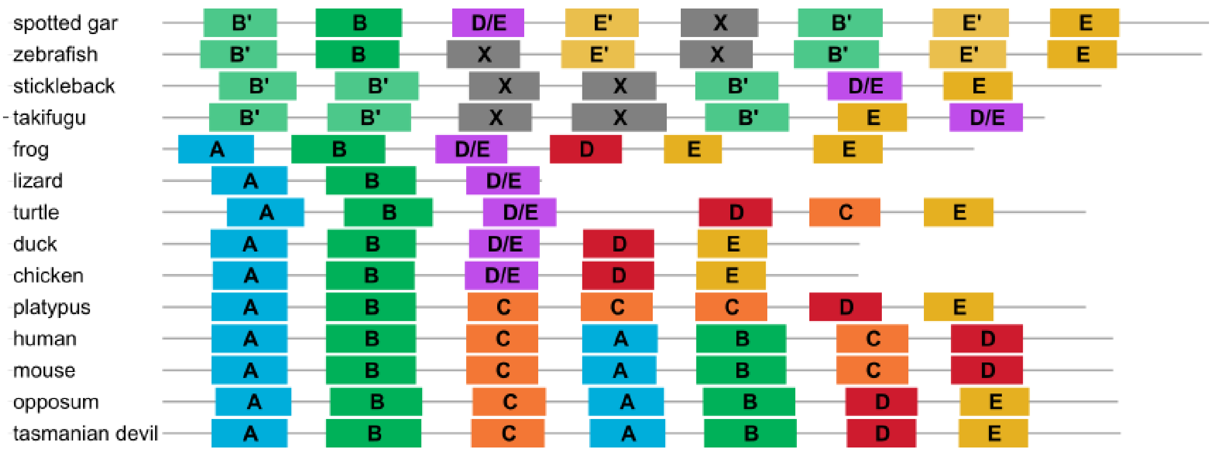


**Figure S6:** *VCAM1 repeat region with repeat units coloured and annotated based on their clustering in the repeat tree and manual inspection, corresponding to the clusters in Figure S5.*

The patterns in sequence similarity of repeat units are used to discern ancestral repeat unit duplications from lineage-specific repeat unit duplications. For example, the most recent common ancestor (MRCA) of tetrapoda already had the A and B repeat units, hence the repeat duplication precedes the speciations. On the other hand, the three adjacent C's in platypus are indicative of a lineage-specific amplification (Figure S7, left). Whilst the pattern in human and mouse hint to a block duplication of A-B-C in the MCRA of theria and putative loss of the second C in the MRCA of opossum and tasmanian devil (metatheria) (Figure S7, left). Another example can be found in the B cluster (Figure S7, right). Here we see indications of two lineage-specific duplications in opossum and tasmanian devil, respectively, and a duplication in the MCRA of human and mouse (eutheria). Note that there is one inconsistent duplication in the repeat tree which is removed during the reconciliation (Figure S7, right).

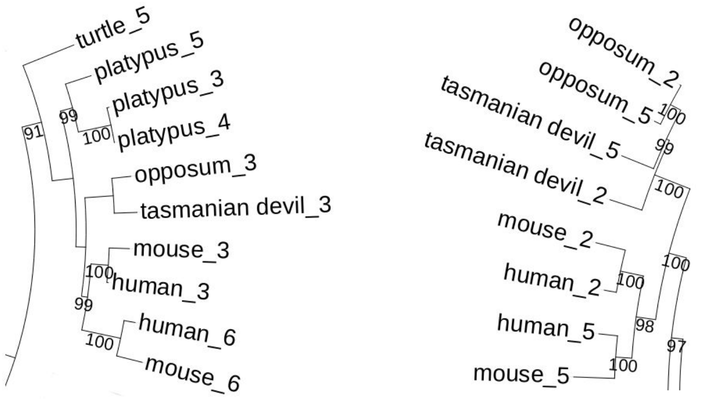


**Figure S7:** *VCAM1 repeat tree detailed view. Left: repeat tree of the "C" cluster of repeat units, right: the "B" repeat units of mammals.*

PhyRepID infers these duplications on their respective gene tree branches consistent with the pattern in the repeat tree (Figure S8). Seven duplications are found in the MCRA of vertebrates and nine dispersed over the gene tree, hence VCAM1 has a moderately fast evolving repeat region compared to other protein families (#242, PRD score 0.48).

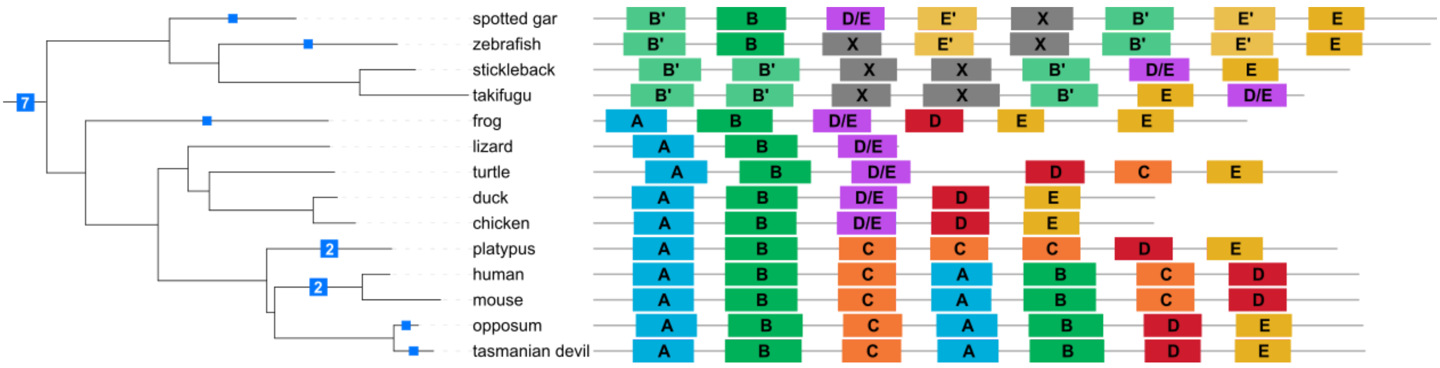


**Figure S8:** *VCAM1 gene tree with duplications as inferred by PhyRepID. The repeat units are coloured and annotated based on their clustering in the repeat tree and manual inspection, corresponding to the clusters in Figure S5.*
