## Supplementary material for "PhyRepID: a comparative phylogenomics approach for large-scale quantification of protein repeat evolution": Supplementary_Note_2_Analysis_results.txt

[1] " ## Dataframe characteristics "
[1] "Total number of OGs: 4939"
[1] "Conserved #OGs <=0 3908, fraction 0.791253290139704"
[1] "Gene dup #OGs >0 3515, fraction 0.711682526827293"
[1] "Human lineage (full) >1 duplication #OGs 1588"
[1] "Mean PRD score: 0.0140689318189107"
[1] "Number of gene trees: 2667"
[1] "Mean PRD score, avg gene tree: 0.0382570083174782"
[1] "## Clan enrichment tests, top 10 clans "
[1] "motif (0.0291152397200673): testing"

	Wilcoxon rank sum test with continuity correction

data:  prd_score by clan_bool
W = 3455498, p-value < 2.2e-16
alternative hypothesis: true location shift is greater than 0

[1] "motif: Fisher enrichment in top 100"

	Fisher's Exact Test for Count Data

data:  fisher.table
p-value = 0.2244
alternative hypothesis: true odds ratio is not equal to 1
95 percent confidence interval:
 0.5112583 1.1776484
sample estimates:
odds ratio 
 0.7766513 

[1] "Beta_propeller (-0.0881163751664286): testing"

	Wilcoxon rank sum test with continuity correction

data:  prd_score by clan_bool
W = 553186, p-value = 9.051e-06
alternative hypothesis: true location shift is less than 0

[1] "Beta_propeller: Fisher enrichment in top 100"

	Fisher's Exact Test for Count Data

data:  fisher.table
p-value = 0.1263
alternative hypothesis: true odds ratio is not equal to 1
95 percent confidence interval:
 0.03978795 1.25427968
sample estimates:
odds ratio 
 0.3348735 

[1] "TPR (-0.10872303800884): testing"

	Wilcoxon rank sum test with continuity correction

data:  prd_score by clan_bool
W = 366790, p-value = 0.0003391
alternative hypothesis: true location shift is less than 0

[1] "TPR: Fisher enrichment in top 100"

	Fisher's Exact Test for Count Data

data:  fisher.table
p-value = 0.0524
alternative hypothesis: true odds ratio is not equal to 1
95 percent confidence interval:
 0.0000000 0.9772754
sample estimates:
odds ratio 
         0 

[1] "C2H2-zf (0.376987284052): testing"

	Wilcoxon rank sum test with continuity correction

data:  prd_score by clan_bool
W = 728016, p-value < 2.2e-16
alternative hypothesis: true location shift is greater than 0

[1] "C2H2-zf: Fisher enrichment in top 100"

	Fisher's Exact Test for Count Data

data:  fisher.table
p-value = 1.474e-15
alternative hypothesis: true odds ratio is not equal to 1
95 percent confidence interval:
  5.561456 14.727891
sample estimates:
odds ratio 
  9.153311 

     [,1] [,2]
[1,]   28  197
[2,]   72 4642
[1] "EF_hand (-0.0819324636797101): testing"

	Wilcoxon rank sum test with continuity correction

data:  prd_score by clan_bool
W = 157384, p-value = 0.1824
alternative hypothesis: true location shift is less than 0

[1] "EF_hand: Fisher enrichment in top 100"

	Fisher's Exact Test for Count Data

data:  fisher.table
p-value = 0.4046
alternative hypothesis: true odds ratio is not equal to 1
95 percent confidence interval:
 0.000000 2.668005
sample estimates:
odds ratio 
         0 

[1] "LRR (-0.118268939412838): testing"

	Wilcoxon rank sum test with continuity correction

data:  prd_score by clan_bool
W = 243468, p-value = 3.545e-11
alternative hypothesis: true location shift is less than 0

[1] "LRR: Fisher enrichment in top 100"

	Fisher's Exact Test for Count Data

data:  fisher.table
p-value = 0.3719
alternative hypothesis: true odds ratio is not equal to 1
95 percent confidence interval:
 0.008031162 1.867336215
sample estimates:
odds ratio 
 0.3224535 

[1] "Ank (-0.0979515279328): testing"

	Wilcoxon rank sum test with continuity correction

data:  prd_score by clan_bool
W = 249538, p-value = 0.0005362
alternative hypothesis: true location shift is less than 0

[1] "Ank: Fisher enrichment in top 100"

	Fisher's Exact Test for Count Data

data:  fisher.table
p-value = 0.5206
alternative hypothesis: true odds ratio is not equal to 1
95 percent confidence interval:
 0.009554968 2.230838594
sample estimates:
odds ratio 
  0.384132 

[1] "Ig (-0.0780670316481818): testing"

	Wilcoxon rank sum test with continuity correction

data:  prd_score by clan_bool
W = 239756, p-value = 0.03987
alternative hypothesis: true location shift is less than 0

[1] "Ig: Fisher enrichment in top 100"

	Fisher's Exact Test for Count Data

data:  fisher.table
p-value = 0.7275
alternative hypothesis: true odds ratio is not equal to 1
95 percent confidence interval:
 0.01089212 2.55219652
sample estimates:
odds ratio 
 0.4383793 

[1] "EGF (-0.0822580796835443): testing"

	Wilcoxon rank sum test with continuity correction

data:  prd_score by clan_bool
W = 154294, p-value = 0.001328
alternative hypothesis: true location shift is less than 0

[1] "EGF: Fisher enrichment in top 100"

	Fisher's Exact Test for Count Data

data:  fisher.table
p-value = 1
alternative hypothesis: true odds ratio is not equal to 1
95 percent confidence interval:
 0.01526502 3.61829267
sample estimates:
odds ratio 
 0.6165966 

[1] "E-set (-0.0390740610267857): testing"

	Wilcoxon rank sum test with continuity correction

data:  prd_score by clan_bool
W = 222169, p-value = 0.0006061
alternative hypothesis: true location shift is less than 0

[1] "E-set: Fisher enrichment in top 100"

	Fisher's Exact Test for Count Data

data:  fisher.table
p-value = 1
alternative hypothesis: true odds ratio is not equal to 1
95 percent confidence interval:
 0.1034741 3.3318028
sample estimates:
odds ratio 
 0.8773867 

[1] "## Categorial enrichment tests  "
[1] "### Schaper "
[1] "Schaper comparison enrichment in top 100 human "
     [,1] [,2]
[1,]    2   90
[2,]   17 1651

	Fisher's Exact Test for Count Data

data:  fisher.table
p-value = 0.2615
alternative hypothesis: true odds ratio is not equal to 1
95 percent confidence interval:
 0.2381323 9.3122262
sample estimates:
odds ratio 
  2.156837 

[1] "Schaper comparison enrichment in  human lin #OGs 1588"
     [,1] [,2]
[1,]   47   45
[2,]  476 1192

	Fisher's Exact Test for Count Data

data:  fisher.table
p-value = 1.072e-05
alternative hypothesis: true odds ratio is not equal to 1
95 percent confidence interval:
 1.674669 4.084420
sample estimates:
odds ratio 
  2.613879 

[1] "Schaper comparison enrichment in human lin #OGs  1588 including unannotated by Schaper as bottom row"
     [,1] [,2]
[1,]   47   45
[2,]  476 1192
[3,] 1065 2114
[1] "Schaper comparison enrichment False in <0 PRD score"
     [,1] [,2]
[1,] 1369  299
[2,]   57   35

	Fisher's Exact Test for Count Data

data:  fisher.table
p-value = 1.354e-05
alternative hypothesis: true odds ratio is not equal to 1
95 percent confidence interval:
 1.755650 4.443836
sample estimates:
odds ratio 
   2.80923 

[1] "Schaper comparison enrichment False in <0 PRD score, including unannotated by Schaper as bottom row"
     [,1] [,2]
[1,] 1369  299
[2,]   57   35
[3,] 2482  697
[1] "### Selectome"
[1] "Selectome enrichment in top 100  PRD score"
     [,1] [,2]
[1,]   12 1063
[2,]   88 3776

	Fisher's Exact Test for Count Data

data:  fisher.table
p-value = 0.01427
alternative hypothesis: true odds ratio is not equal to 1
95 percent confidence interval:
 0.2402399 0.8937732
sample estimates:
odds ratio 
 0.4844695 

[1] "Selectome  comparison enrichment in top 100 human "
     [,1] [,2]
[1,]   20 1055
[2,]   80 3784

	Fisher's Exact Test for Count Data

data:  fisher.table
p-value = 0.7152
alternative hypothesis: true odds ratio is not equal to 1
95 percent confidence interval:
 0.5176481 1.4864946
sample estimates:
odds ratio 
 0.8967022 

[1] "Selectome  comparison enrichment in human lin #OGs 1588"
     [,1] [,2]
[1,]  266  809
[2,] 1322 2542

	Fisher's Exact Test for Count Data

data:  fisher.table
p-value = 2.603e-09
alternative hypothesis: true odds ratio is not equal to 1
95 percent confidence interval:
 0.5400053 0.7388100
sample estimates:
odds ratio 
 0.6322904 

[1] "Comparing PRD score means in Selectome T/F:"
[1] "Selectome True mean: -0.0277789650333953"
[1] "Selectome False mean: 0.0257113979462992"

	Wilcoxon rank sum test with continuity correction

data:  prd_score by selectome
W = 1867154, p-value = 1.826e-07
alternative hypothesis: true location shift is less than 0

[1] "### Segmental duplication regions"
[1] "SegDup enrichment in top 100  PRD score"
     [,1] [,2]
[1,]   20  815
[2,]   80 4024

	Fisher's Exact Test for Count Data

data:  fisher.table
p-value = 0.418
alternative hypothesis: true odds ratio is not equal to 1
95 percent confidence interval:
 0.7118058 2.0486485
sample estimates:
odds ratio 
  1.234299 

[1] "SegDup comparison enrichment in top 100 human "
     [,1] [,2]
[1,]   10  825
[2,]   90 4014

	Fisher's Exact Test for Count Data

data:  fisher.table
p-value = 0.07816
alternative hypothesis: true odds ratio is not equal to 1
95 percent confidence interval:
 0.2496326 1.0467477
sample estimates:
odds ratio 
  0.540666 

[1] "SegDup comparison enrichment in human lin #OGs 1588"
     [,1] [,2]
[1,]  299  536
[2,] 1289 2815

	Fisher's Exact Test for Count Data

data:  fisher.table
p-value = 0.01471
alternative hypothesis: true odds ratio is not equal to 1
95 percent confidence interval:
 1.038294 1.427606
sample estimates:
odds ratio 
  1.218165 

[1] "Comparing PRD score means in SegDup T/F:"
[1] "SegDup True mean: 0.0430799308637126"
[1] "SegDup False mean: 0.00816635282222222"

	Wilcoxon rank sum test with continuity correction

data:  prd_score by segdup
W = 1869106, p-value = 1.616e-05
alternative hypothesis: true location shift is greater than 0

[1] "chi2 independance - Selectome and SegDup"
     [,1] [,2]
[1,]  184  891
[2,]  651 3213

	Pearson's Chi-squared test

data:  tbl
X-squared = 0.043145, df = 1, p-value = 0.8355

[1] "chi2 independance - SegDup and gt>0"
     [,1] [,2]
[1,]  657 2858
[2,]  178 1246

	Pearson's Chi-squared test

data:  tbl
X-squared = 27.653, df = 1, p-value = 1.451e-07

[1] "## EXaC comparison missense z-score and pLI score "
           missense       pLI
dataset   1.2616827 0.4572885
top100    0.6244095 0.2990299
human_lin 1.6492763 0.5334575
[1] "EXac comparision with top 100"

	Wilcoxon rank sum test with continuity correction

data:  mis_z by rank
W = 178039, p-value = 0.003431
alternative hypothesis: true location shift is less than 0


	Wilcoxon rank sum test with continuity correction

data:  pLI by rank
W = 169933, p-value = 0.0004175
alternative hypothesis: true location shift is less than 0

[1] "Exac comparision with human lin #OGs  1588"

	Wilcoxon rank sum test with continuity correction

data:  mis_z by rank_humlin
W = 2665795, p-value < 2.2e-16
alternative hypothesis: true location shift is greater than 0


	Wilcoxon rank sum test with continuity correction

data:  pLI by rank_humlin
W = 2598376, p-value = 2.808e-16
alternative hypothesis: true location shift is greater than 0

[1] "## Gene duplication and oversplit OGs"
[1] "Oversplit OGs (non-euteleostomi root t117571) comparison enrichment in top 100 "
     [,1] [,2]
[1,]   40  446
[2,]   60 4393

	Fisher's Exact Test for Count Data

data:  fisher.table
p-value = 5.974e-16
alternative hypothesis: true odds ratio is not equal to 1
95 percent confidence interval:
  4.232219 10.079910
sample estimates:
odds ratio 
  6.561511 

[1] "Mean gene duplication:"
[1] "dataset: 2.40372545049605"
[1] "top 100: 3.29"
[1] "human_lin: 3.02896725440806"
[1] "Gene duplication T/F enrichment in top 100"
     [,1] [,2]
[1,]   79 3436
[2,]   21 1403

	Fisher's Exact Test for Count Data

data:  fisher.table
p-value = 0.09384
alternative hypothesis: true odds ratio is not equal to 1
95 percent confidence interval:
 0.9351842 2.6281991
sample estimates:
odds ratio 
  1.535981 

[1] "Gene duplication T/F enrichment in human lin #OGs 1588"
     [,1] [,2]
[1,] 1159 2356
[2,]  429  995

	Fisher's Exact Test for Count Data

data:  fisher.table
p-value = 0.05524
alternative hypothesis: true odds ratio is not equal to 1
95 percent confidence interval:
 0.9967048 1.3071266
sample estimates:
odds ratio 
  1.140928 

pdf 
  2 
pdf 
  2 
pdf 
  2 

	Shapiro-Wilk normality test

data:  prd_score
W = 0.17801, p-value < 2.2e-16


	Wilcoxon rank sum test with continuity correction

data:  prd_score by zf_bool
W = 332634, p-value = 1
alternative hypothesis: true location shift is greater than 0


	Wilcoxon rank sum test with continuity correction

data:  prd_score by schaper_pos
W = 97241, p-value = 7.363e-06
alternative hypothesis: true location shift is greater than 0


	Wilcoxon rank sum test with continuity correction

data:  prd_score by gt_bool
W = 3164424, p-value < 2.2e-16
alternative hypothesis: true location shift is not equal to 0


	Wilcoxon rank sum test with continuity correction

data:  prd_score by segdup
W = 1869106, p-value = 1.616e-05
alternative hypothesis: true location shift is greater than 0


	Wilcoxon rank sum test with continuity correction

data:  prd_score by segdup
W = 2983.5, p-value = 0.8332
alternative hypothesis: true location shift is greater than 0


	Wilcoxon rank sum test with continuity correction

data:  prd_score by segdup
W = 543542, p-value = 0.004448
alternative hypothesis: true location shift is greater than 0

null device 
          1 
[1] 486
[1] 4939
